# AlphaFold3 and RoseTTAFold All-Atom structures enable radiosensitizers discovery by targeting multiple DNA damage repair proteins

**DOI:** 10.1101/2025.03.26.645385

**Authors:** Junjie Tong, Xixi Wang, Miaoshan Lu, Tong Wang, Jingying Chen, Jingjing Chen, Yaohan Li, Cong Xie, Yanhui Fu, Changbin Yu

## Abstract

Radioresistance remains a primary obstacle in tumor radiotherapy, with no clinically approved radiosensitizers due to toxicity concerns. To identify effective and safe radiosensitizers, a natural products database containing 79,263 compounds are docked against a hybrid target library of four DNA damage response (DDR)-related proteins, comprising both experimental and artificial intelligence (AI)-predicted structures generated by AlphaFold3 and RoseTTAFold All-Atom models. Retrospectively, AI-modeled structures show comparable *AUC* and *logAUC* values to experimental structures. Prospectively, compounds screened by AI-modeled structures versus those by experimental structures exhibit limited overlap, e.g., 10% for ataxia telangiectasia mutated (ATM), 22.2% for ATM- and Rad3-related (ATR), 7.7% for DNA-dependent protein kinase catalytic subunit (DNA-PKcs), and 40% for Poly (ADP-ribose) polymerase 1 (PARP1). This highlights structural complementarity of AI-modeled structures when docking against small-scale compound libraries. Two compounds exhibiting lower binding free energy than the DNA-PKcs co-crystallized ligand were selected and validated as effective radiosensitizers in tumor cells. Proteomic analyses reveal shared DDR dysregulation but distinct repair pathway vulnerabilities behind both compounds, which activate TP53-associated apoptosis and senescence as cellular endpoints by modulating the synergistic interplay between DDR and spindle checkpoints. These findings highlight their potential as context-dependent radiosensitizers, providing novel candidates and strategies to overcome tumor radioresistance.

## 1. Introduction

Radiotherapy (RT) remains a cornerstone modality in cancer management due to its versatility and efficacy across diverse malignancies. Consecutive exposures to ionizing radiation (IR) can lead to the incidence of radioresistance, a critical factor contributing to RT failure and poor clinical outcomes[1]. DNA damage response (DDR) pathways are the main factors contributing to the development of radioresistance[2]. DNA damage is one of the primary mechanisms by which IR exerts its cytotoxic effects on tumor cells. IR induces lesions, mainly DNA double-strand breaks (DSBs), can lead to cell cycle arrest, apoptosis, or senescence, ultimately resulting in the death of the tumor cells[2]. Progressively, cancer cells have evolved various pathways to repair lesions, including homologous recombination (HR), non-homologous end joining (NHEJ), and alternative end joining(A-EJ). Consequently, targeting key regulators of DNA damage repair pathways has emerged as a strategic approach to restore tumor radiosensitivity[1].

DDR-associated kinases: ataxia telangiectasia mutated (ATM), ATM- and Rad3-related (ATR), and DNA-dependent protein kinase catalytic subunit (DNA-PKcs), and the nuclear enzyme of Poly (ADP-ribose) polymerase 1 (PARP1), represent promising therapeutic targets for cancer treatment[3, 4]. Inhibitors of ATM, ATR, DNA-PKcs, and PARP1 have been developed and demonstrated as effective radiosensitizers for overcoming tumor radioresistance. Currently, ATM inhibitors (AZD0156, AZD1390, M3541, and M4076)[5], ATR inhibitors (M4344, BAY1895344, M6620, AZD6738)[6], DNA-PKcs inhibitors (NVP-BEZ235, LY3023414, CC-115, M9831, M3814, AZD7648)[7] have been investigated in the clinical trials. Moreover, five PARP-1 inhibitors (Talazoparib, Rucaparib, Niraparib, Olaparib, and Veliparib) have been approved by FDA for cancer treatment, but not for use in combination with RT[8]. To date, no clinically approved radiosensitizers exist due to their toxicity concerns[9, 10], highlighting the urgent need for novel agents that balance efficacy with safety. Inhibitors of these DDR regulators have not only demonstrated potential as radiosensitizers, but also induced cancer cell death by their synthetic lethal interactions[10]. Genomic instability, a hallmark of cancer cells often associated with defects in DDR and repair pathways[11], renders cancer cells more susceptible to DNA damage and increasingly dependent on alternative repair mechanisms, thereby underpinning the development of synthetic lethality as a therapeutic strategy in cancer treatment. For instance, ATM deficiency enhances ATR inhibitor efficacy[12, 13], while combined DNA-PKcs and PARP1 inhibitors triggers apoptosis in ATM-deficient tumors[14]. Other interactions include the synergistic effects observed between ATR inhibitors and PARP inhibitors, which can both induce single-strand breaks (SSBs) and exhibit strong synergy in cancer cells[11]. Additionally, DNA-PKcs inhibitors may enhance the efficacy of ATR inhibitor[11]. Similar synthetic lethal interactions have been documented between ATM/DNA-PKcs[15] and ATM/PARP[16], underscoring the therapeutic value of multi-target DDR inhibition. Therefore, targeting DDR regulators of ATM, ATR, DNA-PKcs, and PARP1 may lead to novel radiosensitizers development, or enable cancer treatment by leveraging their multi-target synergistic lethality effects.

Recent advancements in artificial intelligence (AI) have revolutionized structural biology and structure-based drug discovery (SBDD). Following the emergence of AlphaFold2 (AF2)[17], and RoseTTAFold (RF)[18] models, numerous studies were conducted to assess the performance of AI-predicted structures in SBDD. In early retrospective studies, unrefined AF2 structures failed to reproduce ligands screened by experimental structures[19, 20]. While more recently, Lyu et al., discovered that unprocessed AF2 structures prospectively maintained a comparable hit rate to experimental structures and identified novel high-affinity compounds that differ from experimental ones by docking 490 million and 1.6 billion compounds[21]. Compared to the homology model, the AI-predicted structure demonstrated a two-fold increase in the hit rate and identified the most potent agonists when docking a 16 million compound library against a GPCR target[22]. More recently, the development of AlphaFold3 (AF3)[23] and RoseTTAFold All-Atom (RFAA)[24], capable of modeling assemblies of proteins and other biomolecular systems, shows great promise for drug discovery applications. Nevertheless, it remains to be seen whether structures generated by them will deliver enhanced performance in SBDD applications.

In this study, we aimed to discover novel radiosensitizers from a natural products (NP) library by targeting four DDR regulators. NP serves as a major source of lead compounds in drug discovery, with rich structural diversity, high bioactivity, and considerable safety. To date, several NP-sourced radiosensitizers are being under investigation in clinical trials, including papaverine, curcumin, genistein, paclitaxel, and resveratrol[25]. We assembled a hybrid target library containing experimental, AF3-predicted, and RFAA-predicted structures of ATM, ATR, DNA-PKcs, and PARP1. We evaluated the performance of these structures by retrospectively docking them against known ligands and property-match decoys, and prospectively docking the NP subset of ZINC20 database[26]. Compounds identified by cross-docking multiple targets were ranked by molecular dynamics and binding free energy calculations. Two compounds were selected and their anti-proliferative properties and radiosensitization effects upon radiation were validated in tumor cells. Furthermore, their underlying mechanisms were comprehensively characterized though the proteomics analysis, demonstrating their potential as novel DDR-targeting therapeutics.

## 2. Result

### 2.1. Retrospective docking for experimental and AI-modeled structures

We first assessed the difference between experimental and AI-predicted structures, since the performance of AF3 and RFAA models in retrospective docking against ATM, ATR, DNA-PKcs, and PARP1 remains insufficiently evaluated. The experimentally determined structures of ATM (PDB ID: 7NI4, 3.00 Å), ATR (PDB ID: 5YZ0, 4.70 Å), DNA-PKcs (PDB ID: 7OTY, 2.96 Å), and PARP1 (PDB ID: 7AAC, 1.59 Å) were retrieved from the Protein Data Bank (PDB). Structures with superior resolutions and co-bound inhibitors were preferred. While only one experimental structure of ATR has been published to date, and it lacks a co-bound inhibitor. We focused on their primary binding domains: the kinase domains (KD) for ATM, ATR, and DNA-PKcs, and the catalytic domain (CAT) for PARP1. We first evaluated the structural differences between AI-predicted and experimental structures in terms of global structures (Fig.1) and local binding pockets (Figs.1), i.e., residues within 5Å distance from the co-bound ligand. Three AI models (ATM_AF3, PARP1_AF3, and PARP1_RFAA) demonstrated high structural similarity to their experimental counterparts, with global structures and most local residues had root-mean-square deviation (RMSD)<2Å. In contrast, ATR_AF3, ATR_RFAA, and ATM_RFAA had global RMSD>4Å and more local residues with RMSD>2Å.

**Fig. 1:**
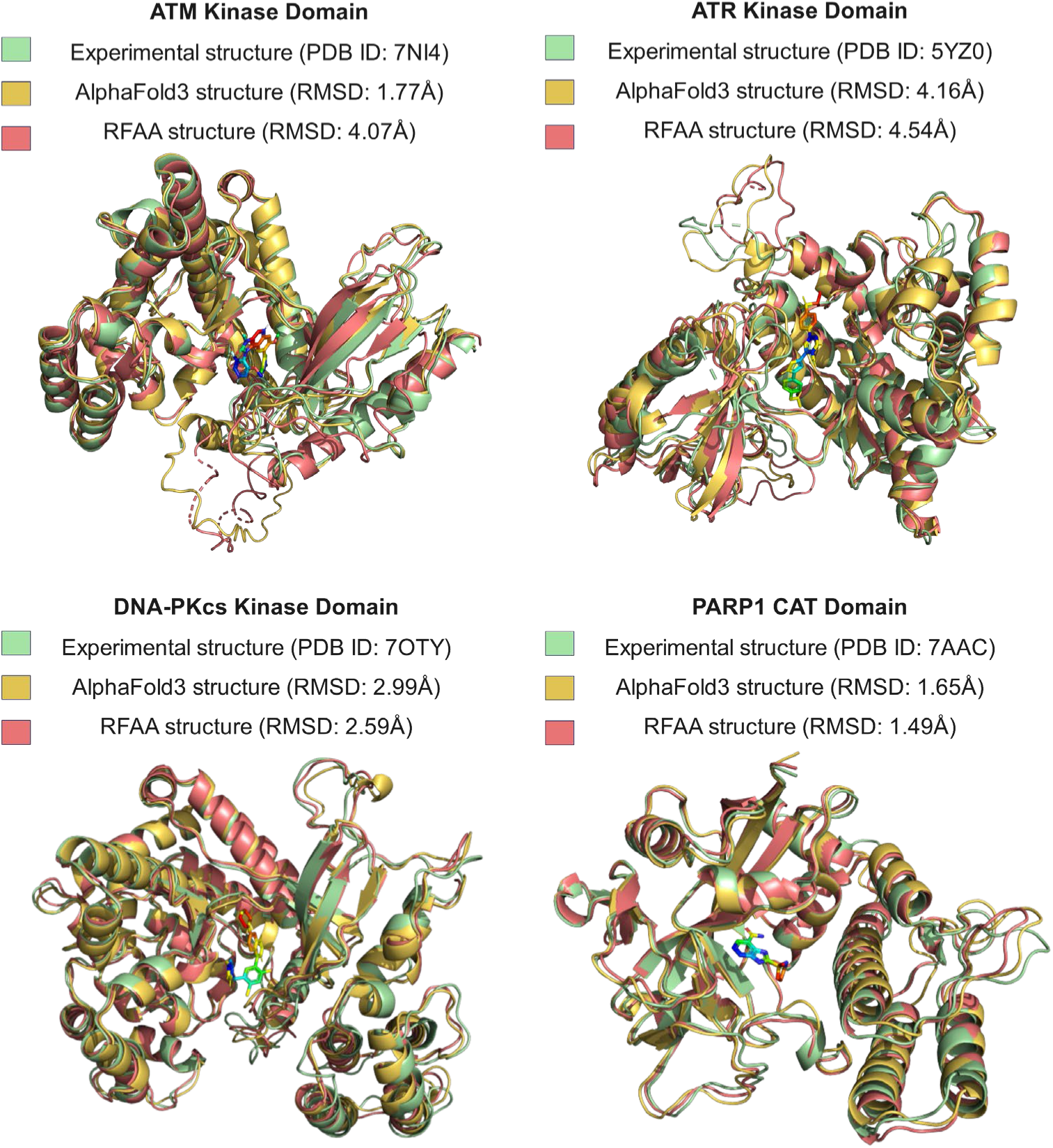
The global differences between experimental and AI-predicted structures for ATM, ATR, DNA-PKcs, and PARP1. Structures were either downloaded from PDB or generated from AF3 and RFAA models. It demonstrated the global RMSD between experimental and AI-modeled structures.

To evaluate the performance of AF3-, RFAA-modeled and experimental structures, we retrospectively docked ATM, ATR, DNA-PKcs, and PARP1 against two types of known ligands: (1) the co-bound inhibitors from experimental assemblies; (2) the known inhibitors retrieved from ChEMBL database[27]. We first docked M4076[28], VX-970[29], M3814[30] and veliparib[31] against AI-predicted and experimental structures of ATM (Figs.2), ATR (Figs.3), DNA-PKcs (Figs.4), and PARP1 (Figs.5), respectively. They accurately predicted the ligands conformations with RMSD < 2Å in all assemblies, determining the docking parameters for the subsequent docking studies.

We prepared known active ligands and their property-matched decoys for each target and docked the active-decoy databases against four structural sources (AF3, RFAA, AF2, and experimental) for each protein (Fig.2). The area under curve (AUC) was calculated to evaluate the overall performance in distinguishing actives from decoys, while the *logAUC* was explored to check their enrichments at the early stages, i.e., 10% top-ranked molecules. For ATM protein (Fig.2A), the experimental (*logAUC* =0.48), RFAA (*logAUC* =0.44), AF2 (*logAUC*=0.49) models showed comparable enrichment, which was higher than the AF3 model (*logAUC* =0.31). In contrast, for the ATR protein (Fig.2B), the experimental structure performed the worst with a *logAUC*=0.21, while the AF3 (*logAUC*=0.55) and AF2 (*logAUC*=0.55) models successfully ranked the active compounds. For DNA-PKcs (Fig. 2C), the AF3 model (*logAUC* = 0.03) exhibited significantly lower enrichment than other models. We calculated the detail parameters of their binding pockets showed that AF3 had the smallest pocket volume (768Å^3^) and the lowest depth (18.73Å), but the highest surface-to-volume ratio (SVR) of 1.34 (Figs.6). For PARP1 (Fig.2D), three structures (AF3, experimental, and AF2) had close *logAUC* values ranged 0.377-0.395, while the RFAA returned a relative lower *logAUC* of 0.272. The previous study demonstrated that crystal structures could double *logAUC* than the AI model retrospectively, but AI-predicted structures successfully identified novel and potent inhibitors in prospective studies[21]. Thus, it suggests that prospective docking campaigns utilizing hybrid target repositories integrating AI-predicted and experimental structures may expand the chemotype of hit compounds in SBDD.

**Fig. 2:**
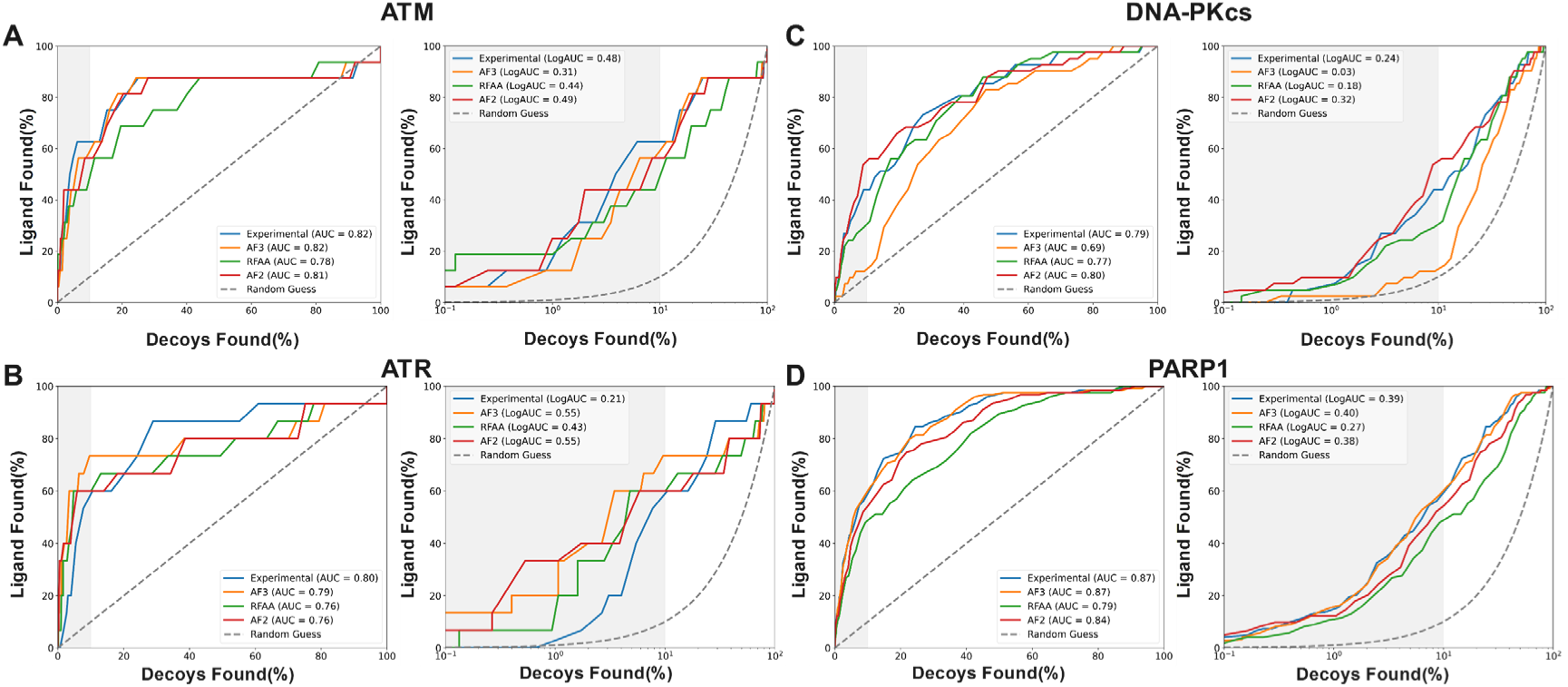
Enrichment analysis of actives against decoys for ATM, ATR, DNA-PKcs, and PARP1. (A) Four structural sources of ATM (Experimental, AF3, AF2, RFAA) were used for retrospective docking screening. The AUC values were used to define the general ability of structures to enrich actives from decoys, while the *logAUC* values described enrichment at early stages. Here, we identified the actives were recognized at top 10% (Gray area) of docked results. The *logAUC* represented the difference between the model’s semilogarthmic AUC and the random semilogarthmic AUC (dashed lines). (B) The AUC and *logAUC* values among different ATR structures. (C) The AUC and *logAUC* values among different DNA-PKcs structures. (D) The AUC and *logAUC* values among different PARP1 structures.

### 2.2. Prospective docking against the Natural Products database

We prospectively docked the AI-experimental hybrid structures against the ZINC20 NP subset, with 79,263 secondary metabolites and its derivatives. We used the affinity score of the 1000th-ranked compound as the cutoff, selecting approximately 1,000 compounds for subsequent evaluation and screening (Table s1). Top-ranked molecules were clustered using Morgan fingerprints, with Tanimoto coefficient (Tc) = 0.5, resulting in clusters numbers ranged from 154 (ATM_RFAA model) to 277 (ATR_AF3 model). Molecules with the highest affinity for each cluster were kept as representative compounds. These representatives were further filtered using stringent druggability and toxicity predictions as described in the Methods section. AI-predicted structures screened eight compounds for ATM (Figs.7), eight for ATR (Figs.8), twelve for DNA-PKcs (Figs.9), and seven for PARP1 (Figs.10). After merging compounds recognized by their corresponding experimental structures, the list of hit compounds expanded: to ten compounds for ATM (overlap ratio of 10%), to nine compounds for ATR (overlap ratio of 22.2%), to thirteen compounds for DNA-PKcs (overlap ratio of 7.7%), to ten compounds for PARP1 (overlap ratio of 40%). This result reflected a low overlap of compounds between AI-predicted and experimental structures. To identify compounds with novel chemotypes, we removed molecules with a Tc value > 0.35 compared to known ligands in the ChEMBL database. We then identified seven potential multi-target compounds: ZINC000001123725 (’3725), ZINC000003797417 (’7417), ZINC000000602850 (’2850), ZINC000000952701 (’2701), ZINC000004085325 (’5325), ZINC000000621966 (’1966), ZINC000004260672 (’0672), by calculating the intersections of four proteins’ candidate lists (Figs.11). These multi-target compounds formed 24 assemblies with different protein structures.

### 2.3. Molecular dynamics simulations and binding free energy calculations of protein-ligand assemblies

To evaluate the binding ability between compounds (′3725, ′7417, ′2850, ′2701, ′5325, ′1966, ′0672) and targets (ATM, ATR, DNA-PKcs, PARP1), we conducted 100 nanoseconds (ns) molecular dynamics simulations (MDS) and the binding free energy (BFE) calculations for all 24 protein-ligand assemblies. Three experimental assemblies, i.e., ATM-M4076, DNA-PKcs-M3814, and PARP1-Veliparib, were considered as positive controls, except for ATR due to its insufficient structural resolution and lack of co-bound ligand in its PDB structure.

We initially performed 100ns MDS on Apo forms of proteins to set the baselines (Figs.12), and experimental assemblies were considered as positive controls. Subsequently, RMSD analyses were conducted: RMSD of the protein backbone was calculated to assess conformational fluctuations, and RMSD of ligands relative to the protein backbone was to evaluate simulation convergence. Additionally, the root-mean-square fluctuation (RMSF) was used to evaluate dynamic characteristics of flexible regions in the proteins. We conducted MDS on five ligands (′2850, ′3725, ′7417, ′2701, and ′5325) formed complexes with ATM (Figs.13), five ligands (′1966, ′0672, ′3725, ′5325, and ′7417) identified by DNA-PKcs (Figs.14), five ligands (′0672, ′1966, ′2701, ′3725, and ′7417) screened by PARP1 (Figs.15), and only two compounds (′2850 and ′3725) filtered by the ATR_AF3 model (Figs.16). The protein backbone RMSDs of assemblies were comparable to their APO forms in most simulation systems, indicating that ligands stabilized their targets structures. Furthermore, most of ligands approached a similar equilibrium state at the end of simulations.

We performed BFE calculations using the last 40ns of the MD trajectory. We used the Molecular mechanics/Poisson–Boltzmann surface area (MM/PBSA) model to calculate the BFE for all complexes and three experimental assemblies, i.e., ATM-M4076 (−28.35±6.7*kcal*/*mol*), DNA-PKcs-M3814 (−30.46± 4.5*kcal*/*mol*), and PARP1-Veliparib (−46.88± 5.0*kcal*/*mol*). Four compounds (′3725, ′7417, ′5325, and ′1966) formed six assemblies, exhibiting improved BFE compared to those reference inhibitors (Table 1). We proceeded to select two compounds (′1966 and ′7417) for further experimental investigation based on their availability for follow-up studies. The ′1966 exhibited stronger binding affinities to both experimental (−46.02±9.4*kcal*/*mol*) and AF3-predicted (−43.03 ± 13.2 *kcal*/*mol*) DNA-PKcs structures than the M3814. While ′1966 also engaged PARP1_PDB, its BFE remained less favorable than the PARP1–veliparib reference system. Similarly, compound ′7417 had a lower BFE than the M3814 when it bound to the DNA-PKcs_RFAA structure (31.35±6.6*kcal*/*mol*). Additionally, ′7417 formed stable interactions with PARP1_PDB (−36.85 ± 7.6 *kcal*/*mol*) and ATM_PDB (−15.45 ± 5.0 *kcal*/*mol*).

**Table 1.** The binding free energy and molecular docking affinity of protein-ligand assemblies. Three reference assemblies were downloaded from PDB database and set as the threshold. Seven potential multi-target compounds formed 24 assemblies with different protein structures.

| NO. | Ligand ID | Protein Name | Structure Source | Binding Free Energy<br>(*kcal/mol*) | Molecular Docking<br>(*kcal/mol*) |
| --- | --- | --- | --- | --- | --- |
| ref | M4076 | ATM | PDB | $-28.35 \pm 6.7$ | -9.5 |
| ref | M3814 | DNA-PKcs | PDB | $-30.46 \pm 4.5$ | -10.4 |
| ref | Veliparib | PARP1 | PDB | $-46.88 \pm 5.0$ | -9.1 |
| 1 | '2850 | ATM | PDB | $-10.52 \pm 8.9$ | -10.2 |
| 2 | '2850 | ATR | AF3 | $-24.26 \pm 5.6$ | -10.9 |
| 3 | '3725 | ATM | PDB | $-33.96 \pm 4.4$ | -10.3 |
| 4 | '3725 | ATM | AF3 | $-19.30 \pm 5.4$ | -10.6 |
| 5 | '3725 | ATR | AF3 | $-27.67 \pm 5.9$ | -11 |
| 6 | '3725 | DNA-PKcs | AF3 | $-21.37 \pm 4.8$ | -10.5 |
| 7 | '3725 | PARP1 | PDB | $-25.81 \pm 8.1$ | -13.6 |
| 8 | '3725 | PARP1 | AF3 | $-38.70 \pm 6.8$ | -13.2 |
| 9 | '3725 | PARP1 | RFAA | $-37.97 \pm 6.2$ | -12.7 |
| 10 | '7417 | ATM | PDB | $-15.45 \pm 5.0$ | -9.0 |
| 11 | '7417 | DNA-PKcs | RFAA | $-31.35 \pm 6.6$ | -10 |
| 12 | '7417 | PARP1 | PDB | $-36.85 \pm 7.6$ | -11.6 |
| 13 | '2701 | ATM | AF3 | $-21.80 \pm 5.4$ | -10.8 |
| 14 | '2701 | PARP1 | PDB | $-21.36 \pm 8.8$ | -12.6 |
| **15** | **'2701** | **PARP1** | **AF3** | **-20.30±8.0** | **-11.8** |
| **16** | **'2701** | **PARP1** | **RFAA** | **-22.64±7.2** | **-11.6** |
| **17** | **'5325** | **ATM** | **RFAA** | **-34.16±12.1** | **-8.7** |
| **18** | **'5325** | **DNA-PKcs** | **AF3** | **-48.31±7.6** | **-9.2** |
| **19** | **'1966** | **DNA-PKcs** | **PDB** | **-46.02±9.4** | **-9.2** |
| **20** | **'1966** | **DNA-PKcs** | **AF3** | **-43.03±13.2** | **-9.2** |
| **21** | **'1966** | **PARP1** | **PDB** | **-18.02±7.7** | **-13.2** |
| **22** | **'1966** | **PARP1** | **RFAA** | **-12.46±8.2** | **-12.4** |
| **23** | **'0672** | **DNA-PKcs** | **AF3** | **-14.75±4.0** | **-9.4** |
| **24** | **'0672** | **PARP1** | **PDB** | **-16.42±5.5** | **-12.4** |

We then analyzed the interaction profiles of ′1966 and ′7417 with their targets at the end of 100ns MDS (Fig.3). Both compounds demonstrated lower BFEs with DNA-PKcs compared to M3814. Notably, ′7417 formed three hydrogen bonds with DNA-PKcs_RFAA at Thr3811 and Ala3730, while ′1966 established stable hydrogen bonds with DNA-PKcs_PDB at Lys3753 and Arg3733, with DNA-PKcs_AF3 at Lys4023 and Lys3753. For ATM and PARP1 targets, the two compounds maintained stable binding within the activate site throughout MDS, but only ′7417 formed a stable hydrogen bond with PARP1_PDB at Glu988, while other complexes were predominantly mediated by hydrophobic interactions.

**Fig. 3:**
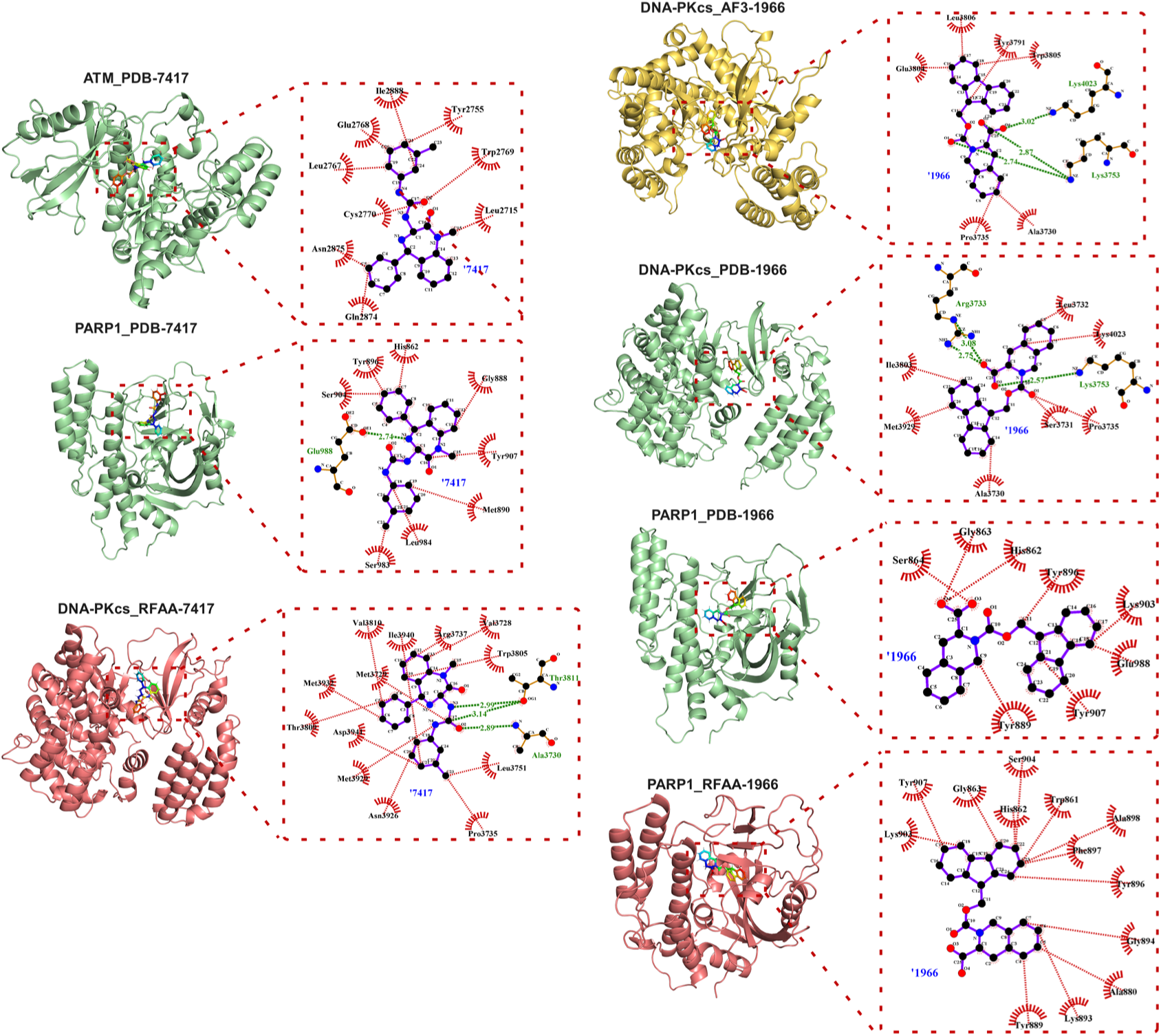
The interaction profiles of ligands (i.e., ′1966 and ′7417) with multiple therapeutic targets. Target structures were derived from AlphaFold3 (AF3) and RoseTTAFold All-Atom (RFAA) predictions and the PDB database, with ligand-target complexes stabilized through 100 ns molecular dynamics (MD) simulations. The left panel demonstrates compound ′7417 engaging with ATM_PDB, PARP1_PDB, and DNA-PKcs_RFAA, while the right panel illustrates compound ′1966 interacting with DNA-PKcs_AF3, DNA-PKcs_PDB, PARP1_PDB, and PARP1_RFAA. Specific interactions were annotated as follows: hydrogen bonds (green dashed lines) and hydrophobic contacts (red dashed lines).

### 2.4. Cellular assays of ′1966 and ′7417

The BFE calculations for ′1966 and ′7417 across multiple targets revealed their multi-target inhibitory potential and candidacy as radiosensitizers (Fig.4A). The ′1966 is a tyrosine derivative featuring a protecting group. It is primarily used as a synthetic intermediate for the synthesis of bioactive peptidomimetics[32]. While the ′7417 compound is a derivative developed and designed from the NP asperlicin[33], demonstrating as a non-peptide gastrin antagonist targeting brain cholecystokinin receptor (CCK-B)[34]. To date, no studies have reported on the relationship between these two compounds and their potential roles as radiosensitizers.

**Fig. 4:**
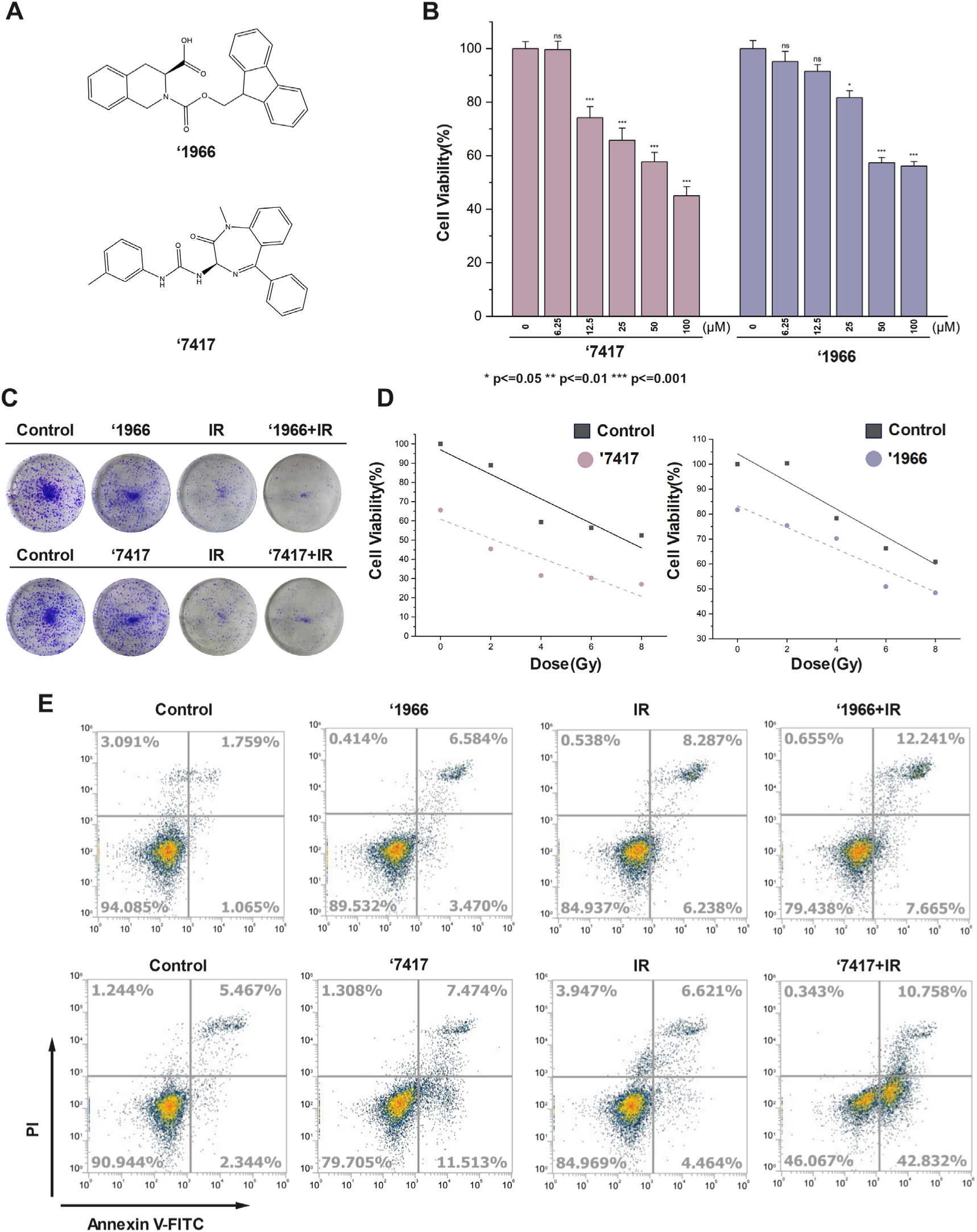
Cell assays of compounds ′1966 and ′7417. (A) The structure of ′1966 and ′7417. (B) A549 cell viability at compounds of ′1966 and ′7417 at concentrations of 6.25µ*M*, 12.5µ*M*, 25µ*M*, 50µ*M*, and 100µ*M*. (C) Colony formation under different conditions. (D) Cell viability under varying radiation doses, i.e., 2*Gy*, 4*Gy*, 6*Gy*, and 8*Gy*, for compound-treated and untreated A549 cells. (E) Flow cytometry was used to detect cells apoptosis under different treatment conditions. The treatment conditions were divided into four groups: normal A549 cells (Control), compound-treated cells, radiation-treated cells (IR), and the combination of compound and radiation-treated cells.

We evaluated their cytostatic activity and radiosensitizing effects upon radiation exposure in A549 cells. Since the synthetic lethality property between DDR proteins, we initially assessed the inhibitory effects of ′1966 and compound ′7417 on A549 cells, revealing that both compounds exhibited dose-dependent suppression of cell viability, with half maximal inhibitory concentration (IC50) > 50 **μM** (Fig.4B). We next investigated their potential roles as radiosensitizer. To maintain cell viability while preserving the inhibitory effects of the compounds, we used 25 **μM** for ′1966 and 12.5 **μM** for ′7417 in subsequent experiments. Upon the additional radiation exposures, we checked the cell viability of compound-treated tumor cells versus untreated cells. The combination of compounds treatment and radiation demonstrated significantly enhanced tumor cell inhibition at doses of 2*Gy*, 4*Gy*, 6*Gy*, and 8*Gy* (Fig.4D). Colony formation assays further confirmed that both compounds, when combined with 4 *Gy* radiation, exhibited superior clonogenic inhibition compared to other groups (Fig.4C). Additionally, flow cytometry assay demonstrated a significant increase in both early and late apoptotic cells in the compound-plus-radiation treatment group (Fig.4E).

### 2.5. Proteomics of ′1966

Compounds ′1966 and ′7417 exhibited validated binding affinities with DDR targets through MDS and BFE calculations, with their anti-proliferative and radiosensitizing effects in tumor cells further establishing their radiosensitizer potential. However, their underlying mechanisms remained unclear, particularly regarding protein-level changes induced by compounds treatment alone or in combination with radiation. To address this, mass spectrometry (MS)-based proteomics was employed to systematically characterize global proteome changes and downstream pathway dysregulation triggered by the compounds, either alone or combined with radiation.

To investigate the mechanistic responses of tumor cells to ′1966, we employed both Data-dependent Acquisition (DDA) and Data-Independent Acquisition (DIA) proteomics across four groups: (1) Untreated tumor cells (Label as “Control”), (2) Cells under radiation (Label as “IR”), (3) Cells treated with compound ′1966(Label as “ZINC1966”), and (4) Cells treated with the compound and radiation (Label as “ZINC1966IR”). We performed Principal Component Analysis (PCA), which effectively distinguished these groups at the proteome level (Fig.5A). We identified 6,645 proteins in the Control group, 6,307 proteins in the ZINC1966 group, 6,380 proteins in the IR group, and 6,261 proteins in the ZINC1966IR group (Fig.5B). The Venn diagram revealed that 5,695 proteins were shared among all four groups (Fig.5C). To explore the differences of compound ′1966 under non-irradiated and irradiated conditions, we extracted the differentially expressed proteins (DEPs), with thresholds of *log2FC*>2, *log2FC*<-2 and *p*<0.05, between:

- the ZINC1966 and Control groups, identifying 205 upregulated proteins and 224 downregulated proteins (Fig.5D).
- the ZINC1966IR and IR groups, identifying 163 upregulated proteins and 310 downregulated proteins (Fig.5E).

**Fig. 5:**
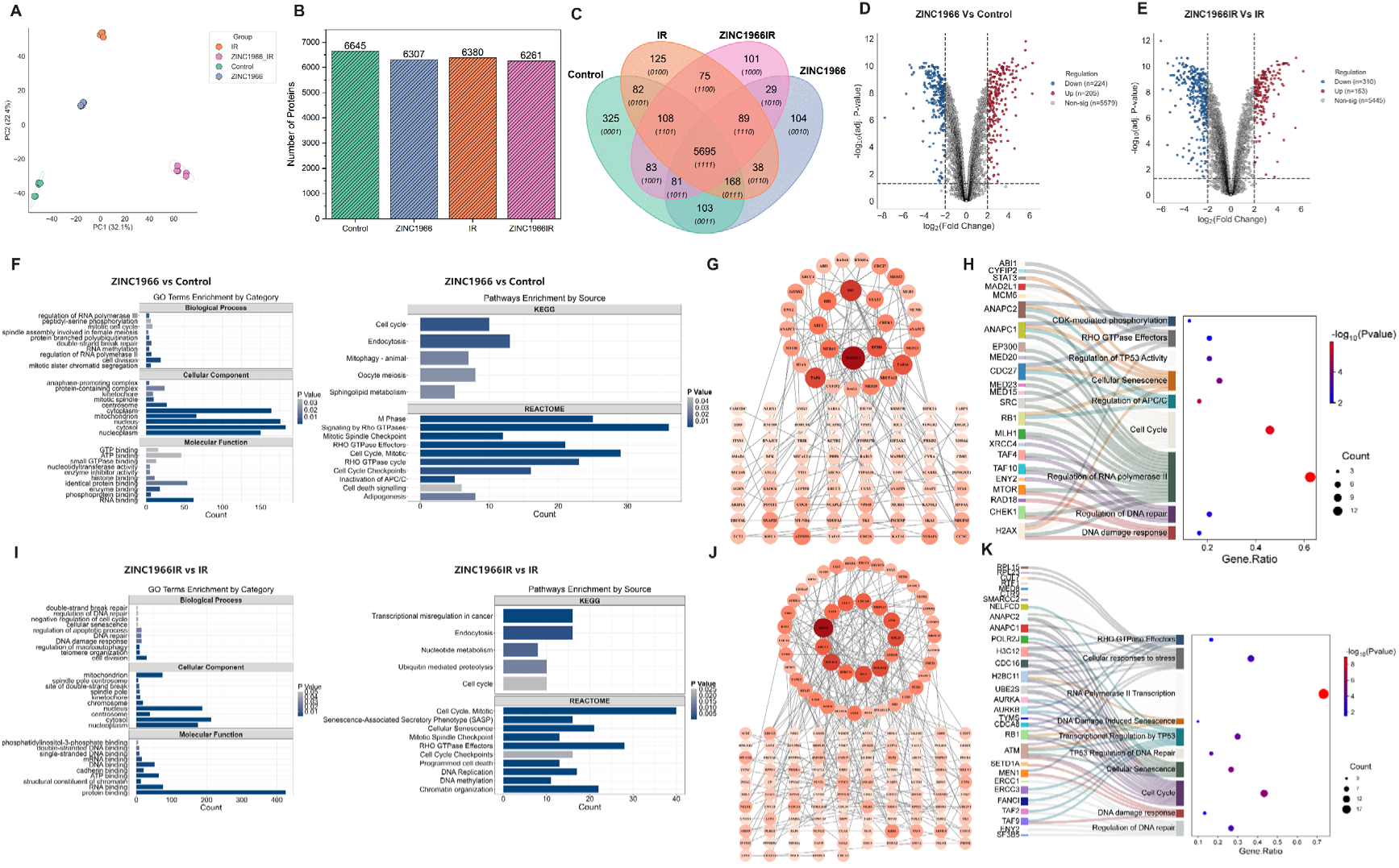
The proteomics analysis of compound 1966. (A) The principal component analysis of four groups (n=5 per group): Control, ZINC1966, IR, and ZINC1966_IR. (B) The number of detected proteins in four groups. (C) The Veen diagram showed overlaps between these four groups. (D) The differentially expressed proteins (DEPs) between ZINC1966 and Control. DEPs were selected based on *log2FC*>2, *log2FC*<-2 and *p*<0.05. (E) The DEPs between ZINC1966IR and IR. (F) Enrichment analysis of DEPs between ZINC1966 and Control using the Gene Ontology (GO), KEGG, and Reactome databases, with *p*<0.05. (G) The protein-protein interactions (PPI) analysis of DEPs between ZINC1966 and Control. These hub genes were identified by the CytoNCA plugin of Cytoscape. (H) Enrichment analysis of hub genes of ZINC1966 vs Control, with *p*<0.05. (I) Enrichment analysis of DEPs between ZINC1966IR and IR using the GO, KEGG, and Reactome databases, with *p*<0.05. (G) The PPI analysis of DEPs between ZINC1966IR and IR. (H) Enrichment analysis of hub genes of ZINC1966IR vs IR, with *p*<0.05.

Next, we performed the enrichment analysis of all DEPs using the Gene Ontology (GO), KEGG, and Reactome databases (Fig.5F). By comparing ZINC1966 against Control, we found that GO-Biological process (BP) was significantly enriched in pathways such as cell division, cell cycle, and double-strand break repair. GO-Cellular component (CC) analysis revealed that the proteins were primarily localized in the nucleoplasm, nucleus, and cytosol. Meanwhile, GO-Molecular function (MF) analysis indicated associations with RNA binding, phosphoprotein binding, and enzyme binding. Further pathway enrichment using KEGG and Reactome identified cell cycle checkpoints, particularly, spindle checkpoints. Subsequently, we conducted Protein-Protein Interaction (PPI) analysis to identify hub genes (Fig.5G), such as CHEK1, EP300, RB1, and MAD2L1. Pathway enrichment of these hub DEPs confirmed significant associations with cell cycle-related pathways, DNA damage response and repair, cellular senescence, and TP53 signaling (Fig.5H).

In the absence of irradiation, the level of DNA damage in cells is limited. To evaluate mechanisms behind the compound under conditions of amplified DNA damage, we analyzed the enrichment results of DEPs between the ZINC1966IR and IR groups (Fig.5I). For the overall DEPs, GO-BP enrichment revealed more associations with DNA damage response, DNA repair (particularly DSB repair), cell cycle regulation, cellular senescence, and apoptosis. GO-CC and GO-MF analyzes indicated that these proteins were primarily localized in the nucleus and sites of cellular damage, with functions closely related to DNA binding. Pathway enrichment analysis identified signaling pathways associated with cellular senescence and programmed cell death as markers of cellular fate, as well as cell cycle checkpoint-related pathways that were downstream of the DNA damage repair. Following PPI analysis (Fig.5J-K), we found that the hub genes remained enriched in pathways related to DNA damage response and repair, cell cycle regulation, TP53-induced apoptosis, and DNA damage-induced cellular senescence.

Based on the PPI analysis of pathways identified by global DEPs and hub genes, we hypothesized that, as a potential radiosensitizer, ′1966 interfered with DNA repair pathway, leading to cell cycle arrest. This ultimately led to TP53-induced apoptosis and cellular senescence as the endpoints of cellular fate. Thus, we annotated the DNA damage response, DDR, Cell cycle checkpoints, Cellular apoptosis, and Cellular senescence pathways by GO_BP, KEGG, Reactome enrichment results of DEPs (Fig.6). Under unirradiated conditions (Fig.6 left panel), we observed the enrichment of DNA damage response-related proteins, such as H2AX (*log2FC*=−7.74), a core protein whose downregulation was previously shown to be associated with increased radiaosensitivity[35]. In DNA repair, by targeting DNA-PKcs and PARP1, part of double-strand and single-strand repair pathways were suppressed, alternative pathways were selectively activated, such as ERCC1(*log2FC* =3.53), which participated in nucleotide excision repair (NER), and MLH1 (*log2FC* =2.34), which was involved in mismatch repair (MMR). The cell cycle checkpoints were primarily enriched in spindle checkpoints. With persistent damage or irreparable DNA damage, the TP53-induced apoptosis and cellular senescence may be triggered. Upon irradiated conditions (Fig.6 right panel), we observed significantly enhanced involvement of the DNA damage response mechanisms, with the activation of alternative DNA repair pathways such as the ATM (*log2FC* =2.25)-mediated pathway, ERCC1 (*log2FC* =4.64)-associated NER, and FANCI (*log2FC*=2.38)-mediated interstrand crosslink (ICL) repair. Similarly, a greater number of cell cycle checkpoints, TP53, apoptosis-, and senescence-related proteins were enriched.

**Fig. 6:**
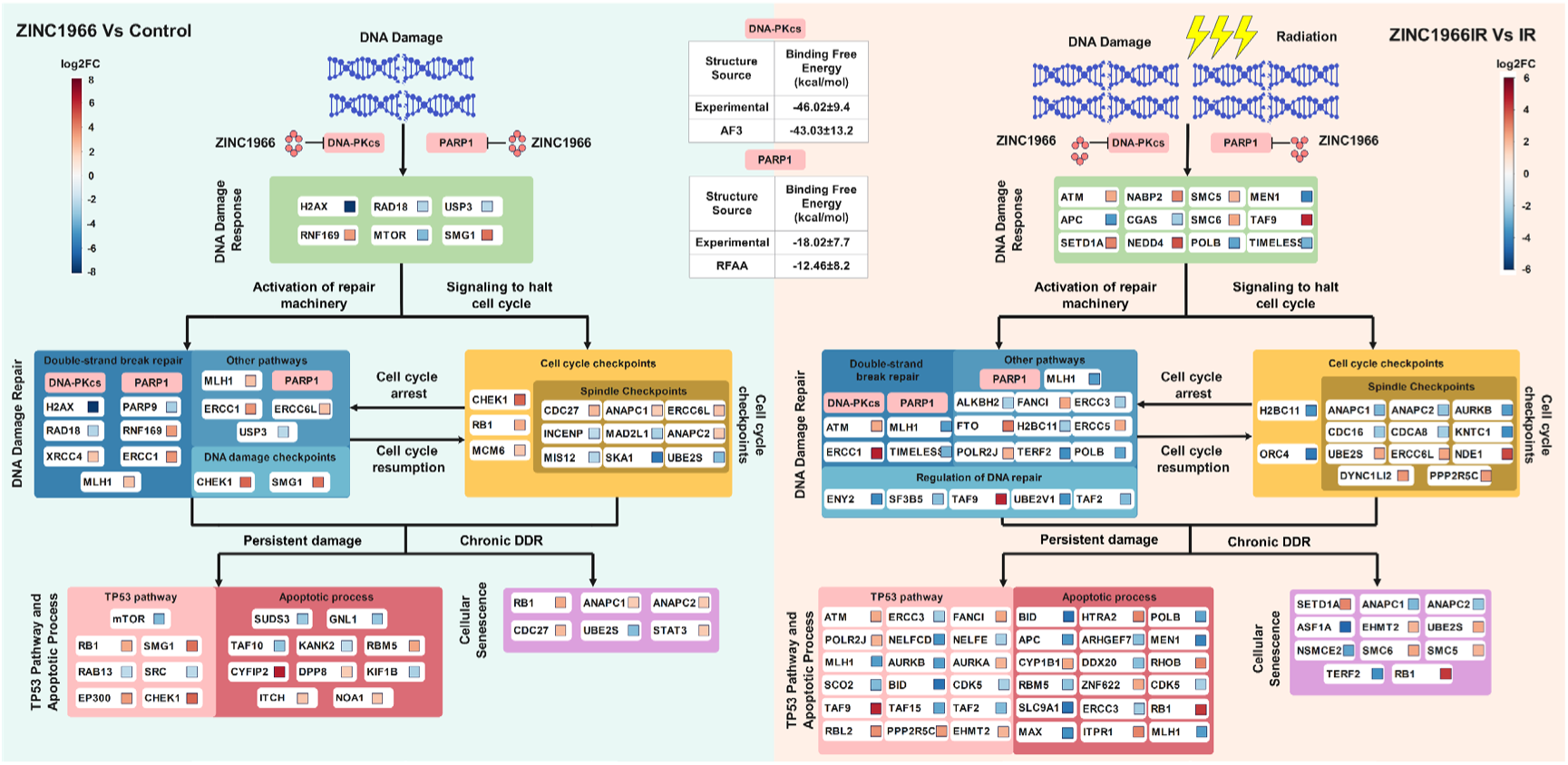
The diagram of the mechanism of compound ′1966 under irradiated and non-irradiated conditions. Interactions of these pathways were identified by EnrichmentMap plugin in Cytoscape. Five pathways were focused on DNA damage response, DNA damage repair, Cell cycle checkpoints, Cell apoptosis and cellular senescence. DEPs were categorized into these pathways and annotated based on *log2FC* values. On the left panel, DEPs between ZINC1966 and Control were annotated. On the right panel, DEPs between ZINC1966IR and IR were annotated.

In summary, our analysis revealed that DDR and cell cycle regulation represented core pathways in both irradiated and unirradiated conditions following treatment with ′1966, while cell apoptosis and senescence acted as downstream endpoints of these cellular responses.

### 2.6. Proteomics of ′7417

Compound ′7417 shared similar targets with ′1966, including DNA-PKcs and PARP1, and the additional secondary target of ATM protein. PCA demonstrated significant differences among four groups: Control, IR, ZINC7417, and ZINC7417IR (Fig.7A). We identified 6,418 proteins in the Control group, 6,362 proteins in the IR group, 6,166 proteins in the ZINC7417 group, and 6,109 proteins in the ZINC7417IR group, with 5,571 proteins shared among all groups (Fig.7B-C). In the DEPs analysis, ZINC7417 exhibited 177 upregulated and 248 downregulated proteins compared to the Control group (Fig.7D). The DEPs between ZINC7417IR and IR groups, we identified 177 upregulated and 213 downregulated proteins (Fig.7E).

**Fig. 7:**
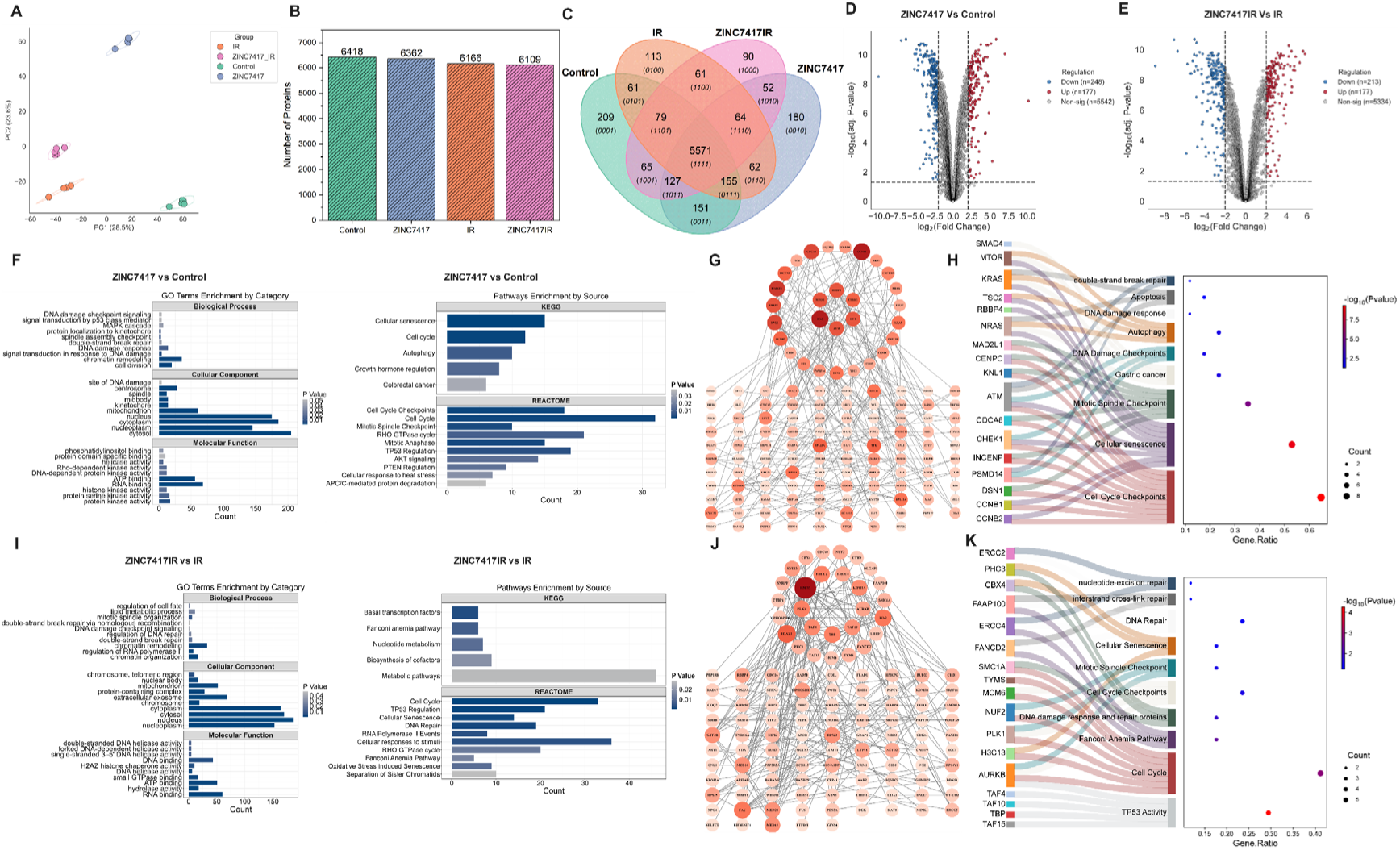
The proteomics analysis of compound 7417. (A) The principal component analysis of four groups (n=5 per group): Control, ZINC7417, IR, and ZINC7417_IR. (B) The number of detected proteins in four groups. (C) The Veen diagram showed overlaps between these four groups. (D) The DEPs between ZINC7417 and Control. DEPs were selected based on *log2FC*>2, *log2FC*<-2 and *p*<0.05. (E) The DEPs between ZINC7417IR and IR. (F) Enrichment analysis of DEPs between ZINC7417 and Control using the GO, KEGG, and Reactome databases, with *p* < 0.05. (G) PPI analysis of DEPs between ZINC7417 and Control. (H) Enrichment analysis of hub genes of ZINC7417 vs Control, with *p* < 0.05. (I) Enrichment analysis of DEPs between ZINC7417IR and IR using the GO, KEGG, and Reactome databases, with *p* < 0.05. (G) The PPI analysis of DEPs between ZINC7417IR and IR. (H) Enrichment analysis of hub genes of ZINC7417IR vs IR, with *p* < 0.05.

In the GO enrichment analysis of DEPs between ZINC7417 and Control (Fig.7F), the BP category included cell cycle pathways and checkpoints, as well as DNA damage response and repair processes. Meanwhile, the MF analysis indicated that the DEPs were predominantly associated with various kinase activities, among which DNA-PKcs and ATM belong to the Ser/Thr protein kinase family. Pathway enrichment analysis identified significant enrichment in cell cycle-related pathways, cellular senescence, autophagy, TP53 pathway, and colorectal cancer-related pathways. PPI analysis identified hub genes related to DDR (Fig.7G), such as ATM, CHEK1, and MTOR. Further pathway enrichment of these hub genes focused on DNA damage response and checkpoint regulation, DSB repair, cell cycle checkpoints, cellular senescence, apoptosis, and autophagy (Fig.7H). Additionally, the gastric cancer pathway was identified, since ′7417, a known CCK-B antagonist, had been associated with gastric cancer and colorectal cancer[33, 34].

Upon additional radiation, we conducted enrichment analysis on the DEPs between ZINC7417IR and IR (Fig.7I). In the GO-BP category, the DEPs were enriched in various DNA damage response and repair-related pathways. Pathway enrichment using KEGG and Reactome demonstrated that DEPs were involved in cellular senescence, cell cycle-related pathways, DNA repair, and Fanconi anemia pathway. Subsequently, we identified hub genes and performed enrichment analysis on them (Fig.7J-K). This revealed enrichment in cell cycle, TP53 activity, cellular senescence, DNA damage response and additional DNA repair pathways, such as NER, ICL repair, mainly Fanconi anemia pathway. Of note, we observed that core enriched pathways were associated with gastric cancer under unirradiated conditions; however, under radiation conditions, we did not observe the enrichment of gastric cancer-related pathways.

Since ′7417 shared similar targets to ′1966, we hypothesized that they exhibit similar mechanisms of action, specifically by regulating DNA repair and cell cycle checkpoints through the DNA damage response, ultimately leading to cell apoptosis and senescence. We next illustrated the mechanism of action for ′7417(Fig.8). Without radiation (Fig.8 left panel), we observed a significant downregulation of ATM (*log2FC* =−2.01) protein, the direct target of ′7417, accompanied by reduced expression of other proteins associated with DSBs, such as XRCC4 (*log2FC* =−3.43). Notably, XRCC4 functions downstream of DNA-PKcs[4]. Consequently, DEPs were significantly enriched in the TP53 pathway and apoptosis and senescence-related pathways. Following radiation exposure (Fig.8 right panel), we did not observe activation of the gastric cancer pathway; however, there were further increases in proteins associated with DNA damage response and repair. Notably, specific repair pathways were activated, such as the Fanconi anemia pathway, including ERCC4 (*log2FC* =2.47), FANCD2 (*log2FC* =3.79), and FAAP100 (*log2FC* =4.53). Additionally, we identified downregulation of numerous proteins involved in negative regulation of apoptotic processes, such as ERCC5 (*log2FC*=−2.63), CBX4 (*log2FC*=−2.04), BABAM2 (*log2FC*=−2.18), MALT1 (*log2FC*=−3.63), and PLK1 (*log2FC*=−3.04).

**Fig. 8:**
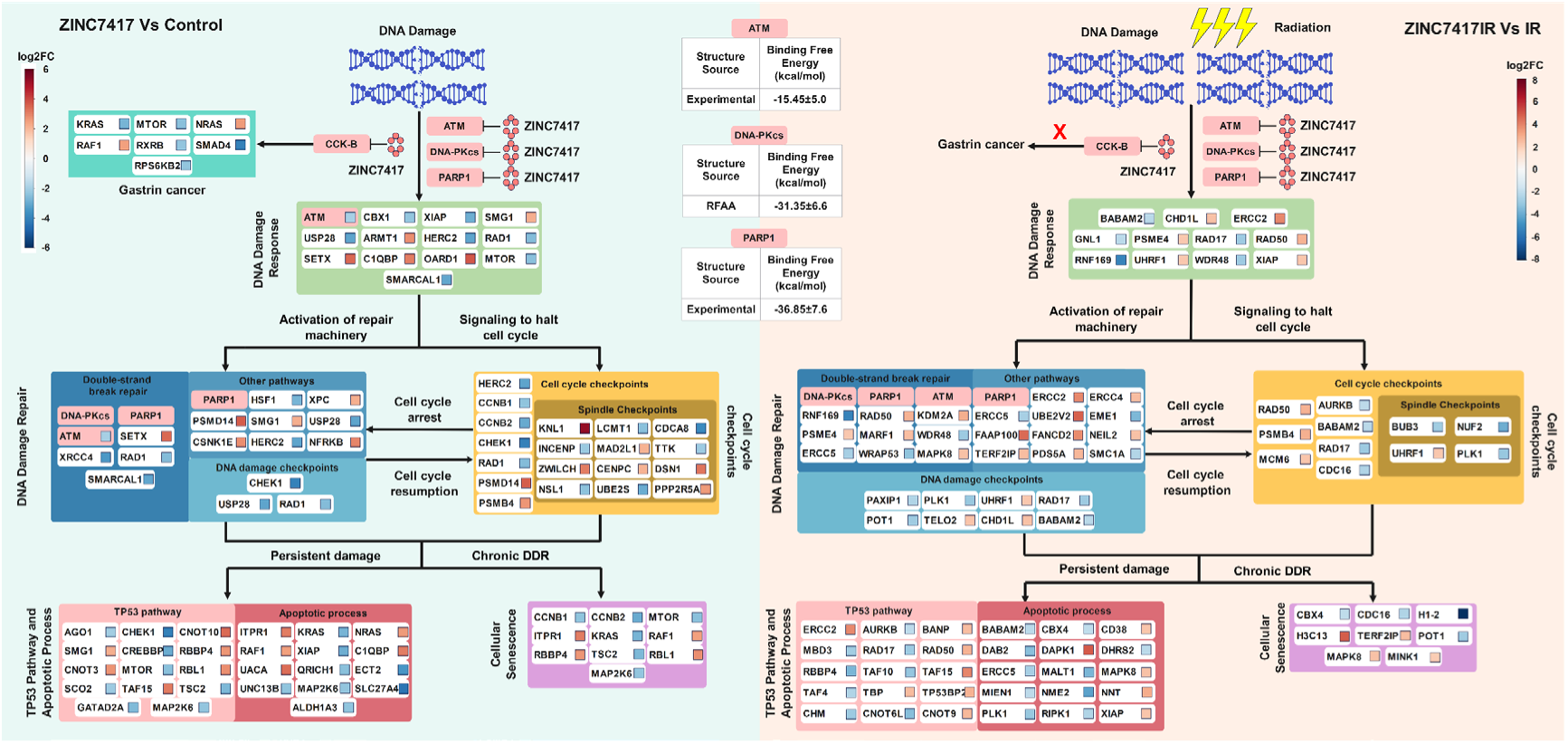
The diagram of the mechanism of compound ′7417 under irradiated and non-irradiated conditions. Interactions of these pathways were identified by EnrichmentMap plugin in Cytoscape. Pathways of Gastrin cancer, DNA damage response, DNA damage repair, Cell cycle checkpoints, Cell apoptosis, and cellular senescence were focused. DEPs were categorized into these pathways and annotated based on *log2FC* values. On the left panel, DEPs between ZINC7417 and Control were annotated in six pathways. On the right panel, DEPs between ZINC7417IR and IR were annotated in five pathways, without the Gastrin cancer.

In summary, the above proteomic analysis demonstrated the potential of both compounds ′1966 and ′7417 as tumor radiosensitizers. Although they are distinct compounds, they exhibit similarities in their mechanisms of action at the proteomic level based on their shared targets.

## 3. Discussion

ATM, ATR, DNA-PKcs, and PARP1, serve as critical components of DDR network and therapeutic targets for developing tumor radiosensitizers. The combined multiple inhibitors of them can effectively exploit the genomic instability of tumor cells to activate synthetic lethality, thereby achieving therapeutic efficacy in cancer treatment and providing a novel strategy to overcome tumor resistance to radiotherapy. Despite the discovery of numerous inhibitors against these targets[5–8], none have been clinically approved for combination with RT, primarily due to dose-limiting toxicity (DLT)[10]. To address this challenge, we performed SBDD on a NP subset from the ZINC20 database to identify safer and more effective radiosensitizers.

We retrospectively and prospectively evaluated the differences on docking performance between AI-predicted and experimental structures, suggesting the applicability of hybrid virtual docking on small-scale library. In retrospective docking campaigns, unlike previous studies[19–21], most AI-predicted structures demonstrated similar or even superior performance in retrospective docking compared to experimental structures. For example, AI-modeled ATR structures exhibited two-fold higher enrichment capacity (*logAUC* 0.43-0.55 vs 0.21 in experimental structure) for active inhibitors (Fig.2), implying that AI modeled better approximation of native protein conformations given the limited resolution (4.70 Å) of the existing PDB structure. These findings suggest that AI models serve as critical alternatives when proteins lack high-quality experimental structures.

In the prospective study, AI-predicted structures screened compounds that differ from experimental structures, which aligns with results from the previous study [21]. The dominance of AI-predicted structures in hit identification was evident across all targets, with AI-screened compounds accounting for eight of ten ATM hits, eight of nine ATR hits, twelve of thirteen DNA-PKcs hits, and seven of ten PARP1 hits (Table S1). Notably, the limited overlap between AI-predicted and experimental structure-identified compounds, range 7.7% for DNA-PKcs to 40% for PARP1, suggests that AI-modeled structures capture distinct chemotype of screened compounds. The discrepancy on compound identification between AI models and experimental conformations was potentially attributable to co-bound ligand-induced binding pocket alterations that restricted compound selection to specific chemotypes. Thus, we propose that the hybrid virtual docking strategy could be a methodology for comprehensive chemical space exploration, circumventing structural biases inherent to single-source experimental conformations, with efficacy in small-scale database screening such as NPs, marketed drugs (thousands of compounds), or even in-house compound collections.

Two novel radiosensitizers, ′1966 and ′7417, were selected by MDS and BFE calculations, and validated in tumor cells. Both compounds exhibited dose-dependent cytotoxic effects in cellular assays (Fig.4A), likely due to synergistic lethal effects from multi-target engagement. We selected compounds ′1966 and ′7417 based on their lower BFE than the reference inhibitor of DNA-PKcs, i.e., M3814 (Table 1). Besides, the ′1966 and ′7417 showed potential interactions with PARP1 and ATM (Table 1). A recent study showed that the DNA-PKcs inhibitor can sensitize cancer cells to IR and, when combined with a PARP inhibitor, can inhibit cell growth and induce apoptosis in ATM-deficient cancer cells[14]. Moreover, this observation has been extended to include interactions between ATM and DNA-PKcs, and between ATM and PARP, elucidating the existence of a synthetic lethal among these key regulators[15, 16]. We observed the inhibition of cancer cells growth by colony formation (Fig.4C) and enhanced apoptosis by flow cytometry (Fig.4E) in compounds treated groups. Furthermore, both compounds demonstrated potential as radiosensitizers, as cell viability decreased further with additional radiation exposure (Fig.4D). These findings provide preliminary evidence supporting the radiosensitizing potential of ′1966 and ′7417.

The cellular phenotypic assays and target-ligand BFEs calculations provide functional readouts and leads, while proteomic profiling quantitatively links observed phenotypes to dysregulated protein network. The proteomic analyses of compounds ′1966 and ′7417, which shared similar targets, revealed distinct yet overlapping mechanisms underlying their roles as potential radiosensitizers. Both compounds modulated DNA damage response and cell cycle checkpoints, with ′1966 primarily interfering with DNA DSB repair pathways (e.g., downregulating H2AX) and activating alternative repair routes (e.g., NER via ERCC1 and MMR via MLH1), while ′7417 exhibited broader kinase inhibition (e.g., ATM) and pathway engagement, including Fanconi anemia-mediated ICL repair. Under non-irradiated conditions, ′1966 induced modest DDR activation and spindle checkpoint enrichment, whereas ′7417 suppressed ATM and downstream DSB repair proteins (e.g., XRCC4), driving TP53-dependent apoptosis and senescence. Radiation amplified these effects: ′1966 enhanced ATM-mediated DDR and alternative repair pathways (e.g., ERCC1, FANCI), while ′7417 upregulated Fanconi anemia components (e.g., FANCD2, ERCC4, FAAP100) and suppressed anti-apoptotic factors (e.g., ERCC5, CBX, BABAM2, MALT1, PLK1), further promoting cell fate endpoints. Notably, ′7417’s association with gastric cancer pathways in non-irradiated conditions—likely linked to its CCK-B antagonism—dissipated upon radiation, suggesting context-dependent therapeutic implications. Both compounds converged on TP53-mediated apoptosis and senescence as terminal outcomes, highlighting DDR and cell cycle dysregulation as central mechanisms. However, their differential engagement of repair pathways (e.g., ATM-mediated repair/NER/MMR for ′1966 vs. Fanconi anemia for ′7417) underscores target-specific vulnerabilities. These findings not only validate their radiosensitizing potential but also emphasize the need for tailored therapeutic strategies based on molecular context, particularly in balancing DDR inhibition and pathway redundancy to optimize tumor-selective cytotoxicity.

Several limitations exist in this study remain to be addressed, such as whether this AI- and experimental hybrid docking strategy remains effective in local laboratory databases containing fewer than 10,000 compounds. While the proteomic analysis of compounds ′1966 and ′7417 demonstrated their impact on DNA repair pathways, further validation of these proteomic insights in preclinical models will clarify their translational relevance. To address these issues, future testing on laboratory-isolated NP databases will be essential. Additionally, more detailed evaluations of radiosensitizers, ′1966 and ′7417, with promising prospects across different tumor types and in vivo efficacy are required in the future.

## 4. Method

### Preparation and assessment of four targets structures

The experimental structures of ATM (PDB ID: 7NI4), ATR (PDB ID: 5YZ0), DNA-PKcs (PDB ID: 7OTY), and PARP1 (PDB ID: 7AAC) were retrieved from the Protein Data Bank (PDB). They formed assemblies with known inhibitors, ATM-M4076, DNA-PKcs-M3814, PARP1-veliparib, except for the ATR which has no bounded inhibitors. We separated the protein structures from ligands and retained the kinase domains (KD) for ATM, ATR, DNA-PKcs, and the catalytic domain (CAT) for PARP1 via PyMol (v2.5.5). The amino acids sequences of key domains of these four proteins were acquired from the UniProt database (ATM ID: Q13315, ATR ID: Q13535, DNA-PKcs ID: P78527, PARP1 ID: P09874). The AF3 structures were generated from the online web tool, AlphaFold Server (https://alphafoldserver.com/). While the RFAA model (https://github.com/baker-laboratory/RoseTTAFold-All-Atom) was deployed on our local servers to predict the protein structures. The root-mean-square deviation (RMSD) of global structures and local residues within 5Å around the binding ligands were visualized and calculated via PyMol and in-house scripts.

### Retrospective molecular docking against known ligands

We initially redocked and docked these co-bound inhibitors to the mixed target library, including 12 experimental and AI-predicted structures of four targets. The molecular docking was performed by QVina2 (https://github.com/QVina/qvina), setting the exhaustiveness value at 12 for ATM and PARP1, at 24 for ATR and DNA-PKcs. Interactions between ligands and targets were further analyzed by PLIP[36] and LigPlot^+^[37].

We prepared the known active ligands from ChEMBL databases[27], with inclusion criteria of affinities (IC50, EC50, Ki, Kd) less than 1μM and molecular weight less than 600*DDkk*. Based on these active ligands, we generated property-matched decoys using the tldr web-service[38]. Thus, we constructed the active-decoys library for target proteins, including 16,972 ligands for ATM, 14,316 ligands for ATR, and 35,540 ligands for DNA-PKcs. Of note, for the PARP1, we retrieved its active-decoys library using the previously prepared one in the DUDE-Z database[39], with 30,543 ligands. We evaluated the ability of each structure to distinguish actives from decoys by calculating the Area Under Curve (AUC) values. Furthermore, we utilized log-adjusted AUC (*logAUC*) to demonstrate the capability of enriching actives with higher rankings. The *logAUC* is defined as:

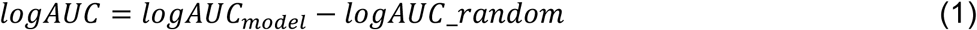

Where equation (1) *logAUC*_*model* is the area under the semilogarithmic ROC curve for the model predictions. Equation (1) *logAUC*_*random* is the area under the semilogarithmic ROC curve for a random guess.

### Prospective molecular docking against the Natural Products Database

In the discovery stage, 79,263 molecules acquired from ZINC20 (https://zinc20.docking.org/) natural products (NP) subset. We prospectively docked the database against the mixed target library of ATM, ATR, DNA-PKcs, and PARP1 using QVina2. We used the affinity score of the 1000th-ranked compound as the cutoff, selecting approximately 1,000 compounds for each structure. These top-ranked compounds were clustered using the 1,024-length binary Morgan fingerprint with a Tanimoto coefficient (Tc)=0.5. Molecules with the highest affinity value in each cluster were retained as the representative compound. The druglikeness, medicinal chemistry property, and pharmacokinetics of representative compounds were predicted using the SwissADME[40]. Five rules from Lipinski’s Rule of Five, Veber’s Rule, Ghose Filter, Muegge Filter, and Egan Filter were applied to check all representative compounds that achieved five “YES” were kept. Then, we kept compounds with predicted High Gastrointestinal (GI) absorption. Besides, the pan-assay interference compounds (PAINS) were excluded. We predicted their AMES toxicity and Carcinogenicity using the ADMETlab 3.0[41]. Compounds with values less than 0.3, labeled as AMES negative and non-carcinogens, were predicted to show excellent safety.

For each target protein, we merged candidates identified by both AI-predict and experimental structures. All candidates were further filtered for novelty by calculating their Morgan fingerprints similarity against corresponding known ligands in the ChEMBL database. Compounds with a Tc value of more than 0.35 were removed from our candidate list.

### Molecular Dynamics Simulations

We performed molecular dynamics simulations (MDS) with GROMACS (version 2024.1)[42]. The protein PDB files were repaired and prepared using PDBFixer[43], including adding missing atoms, residues, and addressing improper formatting. The protonation states(pH=7.4) of ionizable residues in proteins were defined by their pKa (negative logarithm of the acid dissociation constant), which were predicted by PROPKA (v3.5.1)[44]. All ligands were prepared for topology files generation by adding hydrogens(pH=7.4).

The topologies of proteins were prepared by the ′pdb2gmx’ function via the CHARMM36m all-atom force filed[45], and all ligands topology files were generated through the web server of CHARMM General Force Field (CGenFF)[46]. We then rebuilt the protein-ligand complex for following simulations. The complex conformations were placed into a cubic shape box with water molecules, where the distance between the solute and the box edge was set at 1.5nm. To minimize the system, the steepest descent algorithm was used. The minimization process terminated until the maximum force under 1000 kJ mol^−1^nm^−1^ or reached the maximum minimization steps of 50,000. During the equilibration simulations, each protein-ligand complex was position restrained and equilibrated with NVT and NPT ensembles for 100ps, respectively. The MDS runs were performed for 100ns in each complex, and energies were saved per 10ps. The temperature and pressure coupling were controlled by the V-rescale[47] and Parrinello-Rahman[48] methods, with a time constant of 0.1ps and 2ps, reference temperature at 300K and pressure at 1bar, respectively. The short-range van der Waals cutoff was set at 1.2nm and calculated under the Verlet cutoff-scheme. The long-range electrostatics were calculated using Particle Mesh Ewald (PME) method[49]. The hydrogen-bonds were constrained via LINCS algorithm[50]. After the production of MDS runs, the trajectories were used to calculate the RMSD of protein backbones and ligands to evaluate the convergence of the simulation system, and root mean square fluctuation (RMSF) of protein residues to check the flexibility of the protein structure.

### MMPBSA Calculations

The Molecular mechanics/Poisson–Boltzmann (Generalized-Born) surface area (MM/PB(GB)SA) is a popular method to calculate binding free energies (BFE) of protein-ligand complexes. Here, the MMPBSA model was used for our complexes using gmx_MMPBSA(version 1.6.3)[51]. The BFE is normally calculated as the equation below:

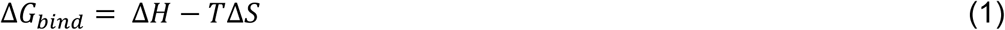

Where equation (1) Δ*H* represents the enthalpy of the binding, while −*T*Δ*S* corresponds to the conformational entropy change upon ligand binding. Since we concerned the relative BFE of ligands towards repeated protein structures, the entropy term was dismissed. The terms of enthalpy are given by:

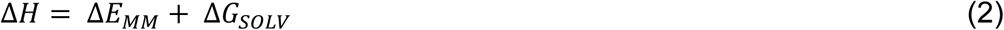

Where equation (2) the gas phase contributions of Δ*E*_*MM*_ and the solvation energy of Δ*G*_*S*OLV_ can be further decomposed into multiple terms:

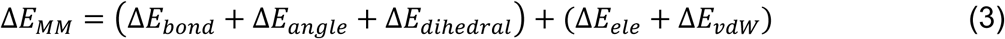

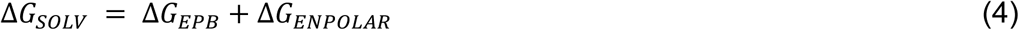

The average BFE calculation for each complex was performed using the last 40ns of trajectory.

### Cell Culture

A549 cells were purchased from Chinese Academy of Sciences Cell Bank (Identifier: SCSP-503) and grown in RPMI1640 media (Gibco) supplemented with 10% fetal bovine serum (Gibco) and 1% penicillin-streptomycin Solution (Beyotime, China). The cell line was incubated at 37℃ with 5% CO_2_ environment (HCP-168, Haier, China).

### Chemicals and Reagents

According to the vendor list on the ZINC20, the compound ′1966, Fmoc-Tic-OH (ZINC ID:621966, purity ≥ 97%, F117108) was purchased from Aladdin Scientific (Shanghai, China). While the compound ′7417, L-365260(ZINC ID:3797417, purity=99.75%, T22895) was purchased from TargetMol Chemicals. During the cell experiments, two compounds were dissolved in dimethyl sulfoxide (DMSO) at 100mM and diluted to 6.25**μM**, 12.5**μM**, 25**μM**, 50**μM** and 100**μM** by the complex medium.

### CCK-8 Assays and Radiation Methods

The cell viability was checked by the CCK-8 assay kit (Beyotime, China). A549 cells were seeded with 5,000 cells per well in the 96-well flat-bottom plates (Beyotime, China) for 24 hours. The medium was then replaced by compounds of ′1966 and ′7417 diluted at five different concentrations. After incubation with ′1966 and ′7417 for another 24 hours, the medium was replaced by 100μL medium with 10μL CCK-8 solution and incubated at 37℃ for 1h. The plates were measured at 450nm using a microplate reader (Spark, Tecan, Switzerland). Together, suitable concentrations of ′1966 and ′7417 were determined by the CCK-8 assays.

To determine the radiation dose for the cell culture, we plated 5,000 cells in each well of the 96-well plates for incubating 24 hours. On the second day, cells were treated by 12.5**μM** ′7417 and 25**μM** ′1966 medium, following by the treatment of radiation doses of 2*Gy*, 4*Gy*, 6*Gy*, and 8*Gy* using the radiator (RS2000PRO, Rad Source Technologies Inc., USA). After another 24 hours incubation, the medium of radiated cells was replaced by CCK-8 medium. The CCK-8 assay was used to evaluate the radiosensitivity of compounds at different radiation dosages.

### Colony formation experiment

The number of 1,000 A549 cells were seeded into each well in 6-well plates. We set up four groups: the control group, the drug-positive group, the IR-positive group, and the drug-IR-positive group. After the cell adhesion was formed, we gave drugs and the 4*Gy* dose of IR to corresponding labeled groups for incubating 24 hours. The medium in each well was replaced by fresh complete medium. After 8 days of incubation, cells were washed with PBS for twice, were fixed by 1 mL 4% paraformaldehyde fix solution (Beyotime, China) for 45 min, and stained by crystal violet (Beyotime, China) for 20 min.

### Flow Cytometry Assay

The cell apoptosis in four groups were measured using the Annexin V-FITC/Propidium Iodide (PI) Apoptosis Detection Kit (Beyotime, China). After treatment of drugs and IR doses, cells were collected and resuspended with the binding buffer, and stained with FITC and PI following the manufacturer instructions. The proportions of apoptotic cells were counted by the flow cytometry (Attune NxT, Thermo Scientific, USA).

### Proteomics Sample Preprocessing

Five bio-replicates were set for cells treated under different conditions. The cell samples were washed by the cold PBS and centrifuged (300g, 5 min, 4℃) for three times. The following steps, including cell lysis, reduction and alkylation, and proteolytic digestion, were processed by the pressure cycling technology (PCT)[52] using the Barocycler 2320EXT (Pressure Biosciences, Inc., USA). After remove the supernatant, 30**μ*L* lysis buffer (8*M* Urea and 2*M* Thiourea, Sigma Aldrich) was added and the sample was transferred into PCT-Micro Tubes (Pressure Biosciences, Inc., USA) for cell lysis (25℃, 60 cycles, 45 kpsi). The reduction and alkylation were conducted by adding 100 *mM* Tris(2-carboxyethyl) phosphine (TCEP, Sigma Aldrich) and 800mM Iodoacetamide (IAA, Sigma Aldrich) stock. Meanwhile, the standard curve of protein concentration was determined using the Pierce BCA Protein Assay Kit (Thermo Scientific, USA) and Nanodrop (Thermo Scientific, USA). The protein amount in experimental samples was calculated by comparing to the standard curve. Two-steps proteolytic digestion was performed with Lys-C (enzyme-to-substrate ratio of 1:40, Hualishi Scientific) for a PCT condition (33℃, 50 cycles, 45), and Trypsin (enzyme-to-substrate ratio of 1:50, Hualishi Scientific) for a PCT condition (33℃, 90 cycles, 45*kpsi*). All reagents used in the PCT steps were dissolved and diluted with 100 *mM* Ammonium bicarbonate (Sigma Aldrich) solution. The digestion reaction was quenched by 20% Trifluoroacetic acid (TFA, Sigma Aldrich) when the *pH* at 2. The peptide desalting procedure was conducted using the hydrophilic-lipophilic balanced (HLB) cartridges (Oasis, Waters), resulting peptides were in 200**μ*L* elution buffer (70% ACN, Sigma Aldrich). The desalted peptides were dried using the SpeedVac SPD1030 (Thermo Scientific, USA) at 35℃, and redissolved in the Buffer (0.1% formic acid (FA), 4% ACN). Finally, the peptide concentrations were measured by Nanodrop and adjusted to 0.1*mg*/**m*L*.

### DDA and DIA Proteomics

A nanoflow DIONEX UltiMate 3000 RSLCnano System connected to an Orbitrap Exploris 480 mass spectrometer equipped with Nanospray Flex ion source and FAIMS Pro Duo system (Thermo Scientific, USA) were used for both data-dependent acquisition (DDA) and data-independent acquisition (DIA) proteomics. Buffer A was H_2_O containing 0.1% FA, and Buffer B was ACN containing 0.1% FA. For the DDA mode, 200*ng* peptides were injected into the LC system and separated by a 120min LC gradient with a flow rate of 300*nL*/*min*. The initial gradient was 3% Buffer B and increased to 8% by 3 min. Buffer B was gradually raised to 35% over 103 min, and reached at 90% in the following 5 min. The column was washed by maintaining a high rate of 90% Buffer B for 5 min. The global parameters of mass spectrometer were set as follows: the FAIMS Mode was standard resolution; the Carrier Gas Flow was at 4.8*L*/*min*; two FAIMS CVs were used, both −65*V* and −45*V*. During the MS1 phase, the scan range of *m*/*z* was from 350 to 1,500 with a resolution of 60,000, the RF lens was set at 50%, the normalized AGC target was 300%, and the maximum injection time was 20*ms*. The time between Masters Scans was set at 3 seconds. Precursors for MS/MS experiments had a minimum signal intensity of 2.0*e*^4^ and charge states from +2 to +6. For MS/MS scans, the isolation window was set at 1.6*m*/*z*, the normalized HCD collision energy was 30%, the resolution was at 15,000, the normalized AGC target was 75%, and the maximum injection time was 22ms. In DIA mode, we used the same LC gradient in DDA proteomics. The parameters for MS1 scans were similar to the DDA experiment, except the scan range was 400-1200*m*/*z*. The 64 variable windows were used in the DIA experiment. The MS/MS spectra were acquired with parameters: the resolution was at 30,000, RF lens was at 50%, the normalized HCD collision energy was 32%, the normalized AGC target was 2,000%, and the maximum injection time was 54*ms*.

### Protein identification and quantification

The resultant DIA MS data were processed by FragPipe(version 22.0)[53] using the DIA_SpecLib_Quant workflow, which builds a spectral library via MSFragger-DIA (version 4.1) and quantifies with DIA-NN (version 1.8.2)[54]. The DIA MS data comprised of two FAIMS CVs (−45 *V* and −65 *V*) were separated into two individual files per CV channel using the FreeStyle (version 1.6, Thermo Scientific), following the recommendations of DIA-NN developers. All DIA and DDA MS data were converted into mzML format files by MSConvert (version 3.0)[55] for the FragePipe workflow. Both DDA and DIA mzML files were used for the spectral library generation. The reviewed human protein database downloaded from Uniport on 25 Sep 2024, containing 40,936 entries (includes 20469 decoys: 50%). The spectral library generated by FragPipe was passed into DIA-NN for the DIA quantification. Of note, the file with only one CV channel was regarded as a separate experiment in DIA-NN, so two CVs reports were combined and merged as one file as described by developers on the DIA-NN Github repository (issue 67).

### Statistical Analysis

During the proteomics data pre-processing, we first established a threshold for missing data, where any protein with more than 50% missing values across all samples in a group was excluded from further analysis. For the remaining proteins with missing values, these were imputed by replacing them with the median value calculated from the non-missing data points. To reduce systematic variation between samples, we applied quantile normalization upon the data using the normalizeBetweenArrays function from the limma package[56] in R software (v4.3.2). Following normalization, the data were *log*2 -transformed to stabilize variance and make the data more normally distributed.

The principal component analysis (PCA) was conducted on the normalized data by Python (v3.9.13). For subsequent differential expression analysis, we then applied linear models and empirical Bayes methods on the data provided by the limma package. Adjusted *p*-values were calculated using the Benjamini-Hochberg method to control the false discovery rate (FDR), and the fold change were *log*2-transformed (*log2FC*). Differentially expressed proteins (DEPs) were identified based on an adjusted p-value threshold of less than 0.05 and a *log2FC*>2 or <-2. The enrichment analysis of DEPs was performed using the DAVID online server[57], annotated by the GO terms, KEGG pathways, and Reactome pathways. The sankey diagram with pathway enrichment dots was plotted by an online platform (https://www.bioinformatics.com.cn).

### Protein-Protein Interaction Analysis

Protein-protein interaction (PPI) network analysis was performed using the STRING database (v12.0)[58] and Cytoscape software (v3.10.2)[59]. We first uploaded DEPs to the STRING platform, with *Homo sapiens* selected as the organism and a high-confidence interaction score threshold set to ≥0.7. Disconnected nodes were excluded to refine the network. The resulting PPI data were exported and imported into Cytoscape for visualization and topological analysis. To identify hub genes, the CytoNCA plugin was employed to select hub genes based on Betweenness (BC), Closeness (CC), and Degree (DC) centralities.

The EnrichmentMap plugin (v3.5.0)[60] in Cytoscape was employed to visualize functional relationships among enriched terms. DAVID output files, enrichment results of DEPs, were directly imported into EnrichmentMap. The edge cutoff was set at 0.375 and the node cutoff was set at *p*-value < 0.05. The results from the EnrichmentMap were subsequently utilized to construct mechanism of action diagrams for the studied drugs.

## Acknowledgements

This work was supported by the Hainan Province Academician Workstation (Changbin Yu), the Specific Research Fund of The Innovation Platform for Academicians of Hainan Province (YSPTZX202407); the Natural Science Foundation of Shandong Province (2022HWYQ-081 and ZR2023LZY009); Scientific Research Cultivation Project in Emerging Strategic Fields (202402); the Academic promotion project of Shandong First Medical University, and funding from Jinan City; the Shandong Center of Technology Innovation for Intelligent Diagnostic; the Mass Spectrometry Big Data Platform of Shandong First Medical University.

## Author contributions

J.T. and C.Y. designed the work; J.T., X.W., M.L. conducted a research and investigation process; J.T., T.W., J.C., J.C. carried out the proteomics experiments; J.T., X.W., M.L. analyzed the data; J.T., X.W., Y.L., C.X., and Y.F. reviewed the manuscript; J.T., M.L., and C.Y. supervised the study; J.T. wrote the paper.

## Competing interest

The authors declare no competing interests.

## Notes

### Competing Interest Statement

The authors have declared no competing interest.

### Summary of Updates

Result section: the interaction profiles of protein-ligands complexes are added. The Introduction and Discussion sections: the language and expression have been rigorously revised to enhance logical coherence, linguistic fluency.

## References

1. Buckley, A.M., et al., Targeting hallmarks of cancer to enhance radiosensitivity in gastrointestinal cancers. Nat Rev Gastroenterol Hepatol, 2020. 17(5): p. 298–313.

2. Huang, R.X. and P.K. Zhou, DNA damage response signaling pathways and targets for radiotherapy sensitization in cancer. Signal Transduct Target Ther, 2020. 5(1): p. 60.

3. Ray Chaudhuri, A. and A. Nussenzweig, The multifaceted roles of PARP1 in DNA repair and chromatin remodelling. Nat Rev Mol Cell Biol, 2017. 18(10): p. 610–621.

4. Blackford, A.N. and S.P. Jackson, ATM, ATR, and DNA-PK: the trinity at the heart of the DNA damage response. Molecular cell, 2017. 66(6): p. 801–817.

5. Du, S., Q. Liang, and J. Shi, Progress of ATM inhibitors: opportunities and challenges. European Journal of Medicinal Chemistry, 2024: p. 116781.

6. Barnieh, F.M., P.M. Loadman, and R.A. Falconer, Progress towards a clinically-successful ATR inhibitor for cancer therapy. Current research in pharmacology and drug discovery, 2021. 2: p. 100017.

7. Matsumoto, Y., Development and evolution of DNA-dependent protein kinase inhibitors toward cancer therapy. International Journal of Molecular Sciences, 2022. 23(8): p. 4264.

8. Cheng, B., et al., Recent advances in DDR (DNA damage response) inhibitors for cancer therapy. European Journal of Medicinal Chemistry, 2022. 230: p. 114109.

9. Yi, J., et al., Potential of natural products as radioprotectors and radiosensitizers: Opportunities and challenges. Food & Function, 2021. 12(12): p. 5204–5218.

10. Drew, Y., F.T. Zenke, and N.J. Curtin, DNA damage response inhibitors in cancer therapy: lessons from the past, current status and future implications. Nature Reviews Drug Discovery, 2025. 24(1): p. 19–39.

11. Hopkins, J.L., L. Lan, and L. Zou, DNA repair defects in cancer and therapeutic opportunities. Genes & development, 2022. 36(5-6): p. 278–293.

12. Villaruz, L.C., et al., ATM protein is deficient in over 40% of lung adenocarcinomas. Oncotarget, 2016. 7(36): p. 57714.

13. Liu, T.-T., et al., Discovery of a meisoindigo-derived protac as the ATM degrader: Revolutionizing colorectal cancer therapy via synthetic lethality with ATR inhibitors. Journal of Medicinal Chemistry, 2024. 67(9): p. 7620–7634.

14. Fok, J.H., et al., AZD7648 is a potent and selective DNA-PK inhibitor that enhances radiation, chemotherapy and olaparib activity. Nature communications, 2019. 10(1): p. 5065.

15. Gurley, K.E. and C.J. Kemp, Synthetic lethality between mutation in Atm and DNA-PKcs during murine embryogenesis. Current Biology, 2001. 11(3): p. 191–194.

16. Wang, C., et al., Genetic vulnerabilities upon inhibition of DNA damage response. Nucleic acids research, 2021. 49(14): p. 8214–8231.

17. Jumper, J., et al., *Highly accurate protein structure prediction with AlphaFold*. nature, 2021. 596(7873): p. 583–589.

18. Baek, M., et al., Accurate prediction of protein structures and interactions using a three-track neural network. Science, 2021. 373(6557): p. 871–876.

19. Díaz-Rovira, A.M., et al., *Are deep learning structural models sufficiently accurate for virtual screening? Application of docking algorithms to AlphaFold2* predicted structures. Journal of Chemical Information and Modeling, 2023. 63(6): p. 1668–1674.

20. Karelina, M., J.J. Noh, and R.O. Dror, How accurately can one predict drug binding modes using AlphaFold models? Elife, 2023. 12: p. RP89386.

21. Lyu, J., et al., AlphaFold2 structures guide prospective ligand discovery. Science, 2024. 384(6702): p. eadn6354.

22. Díaz-Holguín, A., et al., AlphaFold accelerated discovery of psychotropic agonists targeting the trace amine–associated receptor 1. Science Advances, 2024. 10(32): p. eadn1524.

23. Abramson, J., et al., Accurate structure prediction of biomolecular interactions with AlphaFold 3. Nature, 2024. 630(8016): p. 493–500.

24. Krishna, R., et al., Generalized biomolecular modeling and design with RoseTTAFold All-Atom. Science, 2024. 384(6693): p. eadl2528.

25. Komorowska, D., et al., Natural radiosensitizers in radiotherapy: Cancer treatment by combining ionizing radiation with resveratrol. International Journal of Molecular Sciences, 2022. 23(18): p. 10627.

26. Irwin, J.J., et al., ZINC20—a free ultralarge-scale chemical database for ligand discovery. Journal of chemical information and modeling, 2020. 60(12): p. 6065–6073.

27. Zdrazil, B., et al., The ChEMBL Database in 2023: a drug discovery platform spanning multiple bioactivity data types and time periods. Nucleic acids research, 2024. 52(D1): p. D1180–D1192.

28. Fuchss, T., et al., Highly potent and selective ATM kinase inhibitor M4076: A clinical candidate drug with strong anti-tumor activity in combination therapies. Cancer Research, 2019. 79(13_Supplement): p. 3500–3500.

29. Yap, T.A., et al., Phase I trial of first-in-class ATR inhibitor M6620 (VX-970) as monotherapy or in combination with carboplatin in patients with advanced solid tumors. Journal of Clinical Oncology, 2020. 38(27): p. 3195–3204.

30. van Bussel, M.T., et al., A first-in-man phase 1 study of the DNA-dependent protein kinase inhibitor peposertib (formerly M3814) in patients with advanced solid tumours. British Journal of Cancer, 2021. 124(4): p. 728–735.

31. Coleman, R.L., et al., Veliparib with first-line chemotherapy and as maintenance therapy in ovarian cancer. New England Journal of Medicine, 2019. 381(25): p. 2403–2415.

32. Kumar, V. and J.V. Aldrich, A solid-phase synthetic strategy for labeled peptides: synthesis of a biotinylated derivative of the δ opioid receptor antagonist TIPP (Tyr-Tic-Phe-Phe-OH). Organic Letters, 2003. 5(5): p. 613–616.

33. Bock, M.G., et al., Benzodiazepine gastrin and brain cholecystokinin receptor ligands; L-365,260. Journal of medicinal chemistry, 1989. 32(1): p. 13–16.

34. Lotti, V.J. and R.S. Chang, A new potent and selective non-peptide gastrin antagonist and brain cholecystokinin receptor (CCK-B) ligand: L-365,260. European journal of pharmacology, 1989. 162(2): p. 273–280.

35. Yang, Z., et al., miR-328-3p enhances the radiosensitivity of osteosarcoma and regulates apoptosis and cell viability via H2AX. Oncology reports, 2018. 39(2): p. 545–553.

36. Adasme, M.F., et al., PLIP 2021: expanding the scope of the protein–ligand interaction profiler to DNA and RNA. Nucleic acids research, 2021. 49(W1): p. W530–W534.

37. Laskowski, R.A. and M.B. Swindells, LigPlot+: multiple ligand–protein interaction diagrams for drug discovery. 2011, ACS Publications.

38. Irwin, J.J., et al., Automated docking screens: a feasibility study. Journal of medicinal chemistry, 2009. 52(18): p. 5712–5720.

39. Stein, R.M., et al., Property-unmatched decoys in docking benchmarks. Journal of chemical information and modeling, 2021. 61(2): p. 699–714.

40. Daina, A., O. Michielin, and V. Zoete, SwissADME: a free web tool to evaluate pharmacokinetics, drug-likeness and medicinal chemistry friendliness of small molecules. Scientific reports, 2017. 7(1): p. 42717.

41. Fu, L., et al., ADMETlab 3.0: an updated comprehensive online ADMET prediction platform enhanced with broader coverage, improved performance, API functionality and decision support. Nucleic acids research, 2024. 52(W1): p. W422–W431.

42. Abraham, M.J., et al., GROMACS: High performance molecular simulations through multi-level parallelism from laptops to supercomputers. SoftwareX, 2015. 1: p. 19–25.

43. Eastman, P., et al., OpenMM 7: Rapid development of high performance algorithms for molecular dynamics. PLoS computational biology, 2017. 13(7): p. e1005659.

44. Olsson, M.H., et al., PROPKA3: consistent treatment of internal and surface residues in empirical p K a predictions. Journal of chemical theory and computation, 2011. 7(2): p. 525–537.

45. Huang, J., et al., CHARMM36m: an improved force field for folded and intrinsically disordered proteins. Nature methods, 2017. 14(1): p. 71–73.

46. Vanommeslaeghe, K., et al., CHARMM general force field: A force field for drug-like molecules compatible with the CHARMM all-atom additive biological force fields. Journal of computational chemistry, 2010. 31(4): p. 671–690.

47. Berendsen, H.J., et al., Molecular dynamics with coupling to an external bath. The Journal of chemical physics, 1984. 81(8): p. 3684–3690.

48. Parrinello, M. and A. Rahman, Polymorphic transitions in single crystals: A new molecular dynamics method. Journal of Applied physics, 1981. 52(12): p. 7182–7190.

49. Darden, T., D. York, and L. Pedersen, Particle mesh Ewald: An N log (N) method for Ewald sums in large systems. Journal of chemical physics, 1993. 98: p. 10089–10089.

50. Hess, B., et al., LINCS: A linear constraint solver for molecular simulations. Journal of computational chemistry, 1997. 18(12): p. 1463–1472.

51. Valdés-Tresanco, M.S., et al., gmx_MMPBSA: a new tool to perform end-state free energy calculations with GROMACS. Journal of chemical theory and computation, 2021. 17(10): p. 6281–6291.

52. Cai, X., et al., High-throughput proteomic sample preparation using pressure cycling technology. Nature Protocols, 2022. 17(10): p. 2307–2325.

53. Yu, F., et al., Analysis of DIA proteomics data using MSFragger-DIA and FragPipe computational platform. Nature communications, 2023. 14(1): p. 4154.

54. Demichev, V., et al., DIA-NN: neural networks and interference correction enable deep proteome coverage in high throughput. Nature methods, 2020. 17(1): p. 41–44.

55. Chambers, M.C., et al., A cross-platform toolkit for mass spectrometry and proteomics. Nature biotechnology, 2012. 30(10): p. 918–920.

56. Ritchie, M.E., et al., *limma powers differential expression analyses for RNA-sequencing and microarray studies*. Nucleic acids research, 2015. 43(7): p. e47–e47.

57. Sherman, B.T., et al., DAVID: a web server for functional enrichment analysis and functional annotation of gene lists (2021 update). Nucleic acids research, 2022. 50(W1): p. W216–W221.

58. Szklarczyk, D., et al., The STRING database in 2021: customizable protein–protein networks, and functional characterization of user-uploaded gene/measurement sets. Nucleic acids research, 2021. 49(D1): p. D605–D612.

59. Shannon, P., et al., Cytoscape: a software environment for integrated models of biomolecular interaction networks. Genome research, 2003. 13(11): p. 2498–2504.

60. Merico, D., et al., Enrichment map: a network-based method for gene-set enrichment visualization and interpretation. PloS one, 2010. 5(11): p. e13984.

