## Supplemental figures and tables for "AlphaFold3 and RoseTTAFold All-Atom structures enable radiosensitizers discovery by targeting multiple DNA damage repair proteins"

### Table of contents


**Figs.1:** The differences of binding pockets between experimental and AI- predicted structures for ATM, ATR, DNA-PKcs, and PARP1. ......................................................S3

**Figs.2:** The interaction analysis of ligand M4076 docked with the experimental structure, AF3, and RFAA model of ATM. ..........................................................................S4

**Figs.3:** The interaction analysis of ligand VX970 docked with the experimental structure, AF3, and RFAA model of ATR. . .........................................................................S5

**Figs.4:** The interaction analysis of ligand M3814 docked with the experimental structure, AF3, and RFAA model of DNA-PKcs. . .............................................................S6

**Figs.5:** The interaction analysis of ligand veliparib docked with the experimental structure, AF3, and RFAA model of PARP1. . ..................................................................S7

**Figs.6:** The binding pockets parameters of DNA-PKcs structrues. . ..............................S8

**Figs.7:** The filtered compounds identified by ATM structures. . .....................................S9

**Figs.8:** The filtered compounds identified by ATR structures. . ...................................S10

**Figs.9:** The filtered compounds identified by DNA-PKcs structures. . ..........................S11

**Figs.10:** The filtered compounds identified by PARP1 structures. ..............................S12

**Figs.11:** Seven potential multi-target compounds. . ....................................................S13

**Figs.12:** The molecular dynamics simulations (MDS) of Apo forms of 12 protein structures. ...................................................................................................................S14

**Figs.13:** The 100ns molecular dynamics simulations (MDS) of protein-ligand assemblies screened by ATM protein. ...........................................................................................S15

**Figs.14:** The 100ns molecular dynamics simulations (MDS) of protein-ligand assemblies screened by DNA-PKcs protein. ......................................................................S16

**Figs.15:** The 100ns molecular dynamics simulations (MDS) of protein-ligand assemblies screened by PARP1 protein. ...........................................................................S17

**Figs.16:** The 100ns molecular dynamics simulations (MDS) of protein-ligand assemblies screened by ATR structure. ..........................................................................S18

**Table s1**: Changes in the number of compounds in target structures with different filtering criteria. ..............................................................................................S19

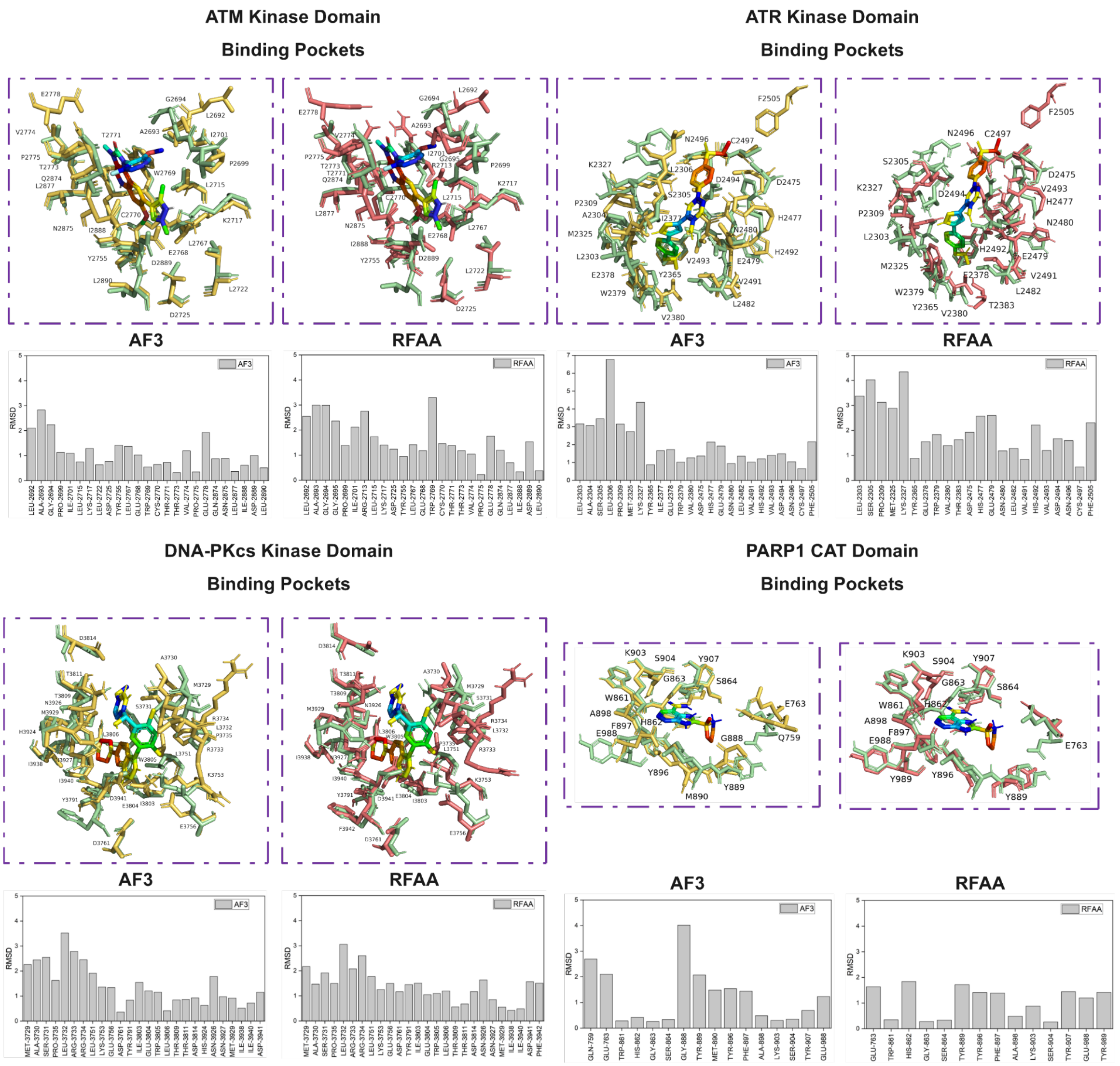


Figs. 1: The differences of binding pockets between experimental and AI-predicted structures for ATM, ATR, DNA-PKcs, and PARP1. Structures were either downloaded from PDB or generated from AF3 and RFAA models. It showed the RMSD of residues within 5Å distance from the co-bound inhibitors were calculated for local difference evaluations.

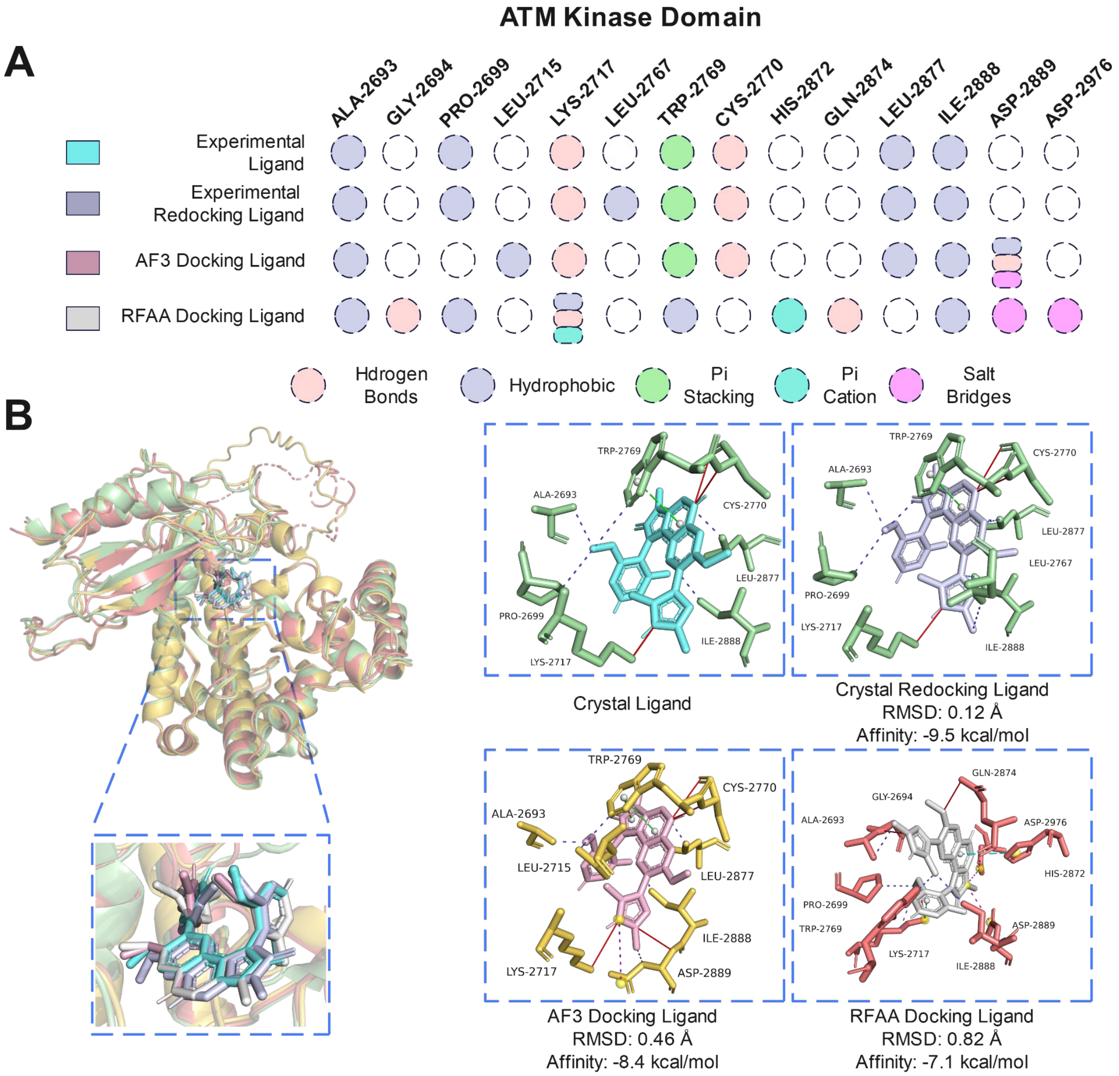


Figs. 2: The interaction analysis of ligand M4076 docked with the experimental structure, AF3, and RFAA model of ATM. (A) The interactions between M4076 and ATM KD structures from different sources. (B) Conformations of M4076 with experimental and AI-predicted ATM KD structures, along with the RMSD values compared to the original structure and the affinity scores from molecular docking.

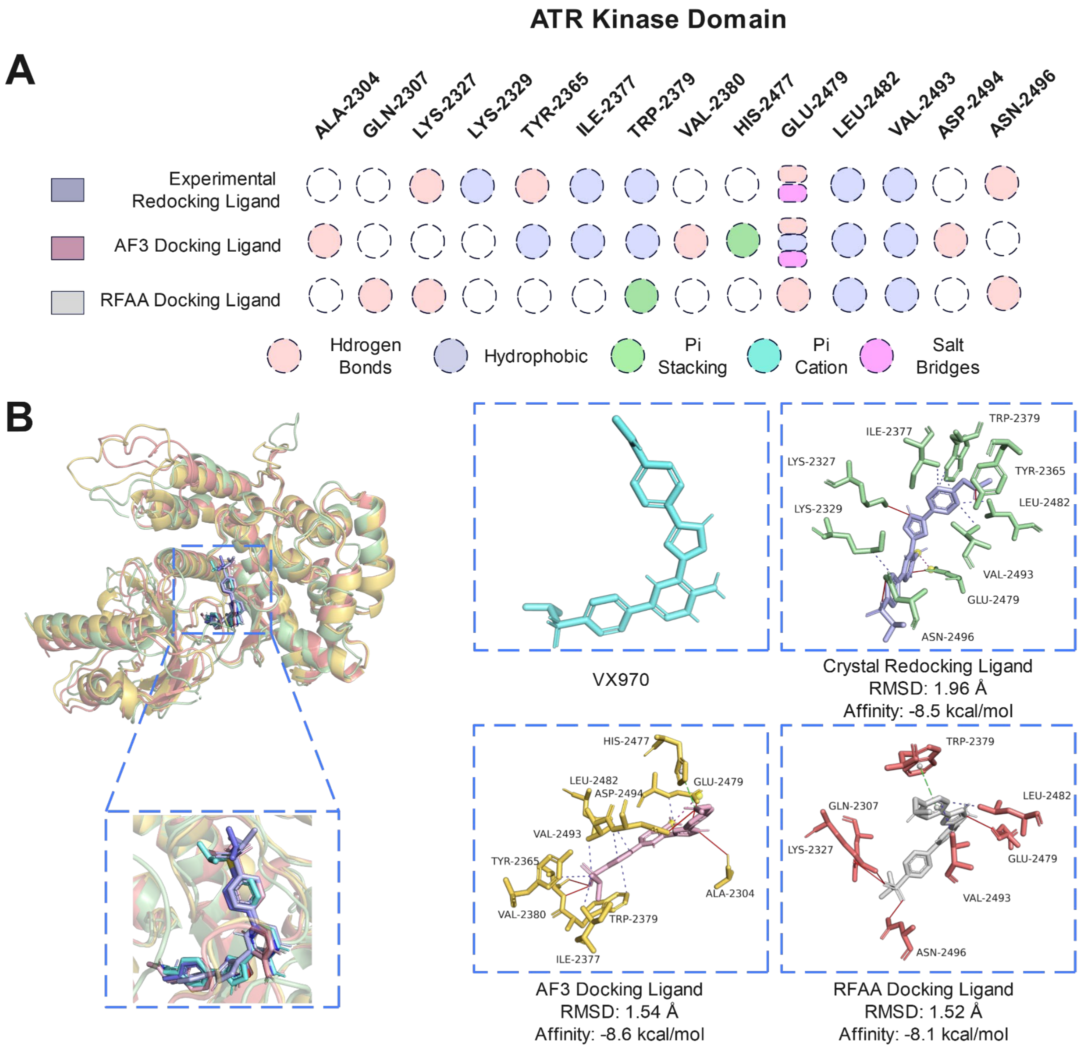


Figs. 3: The interaction analysis of ligand VX970 docked with the experimental structure, AF3, and RFAA model of ATR. (A) The interactions between VX970 and ATR KD structures from different sources. (B) Conformations of VX970 with experimental and AI-predicted ATR KD structures, along with the RMSD values compared to the original structure and the affinity scores from molecular docking.

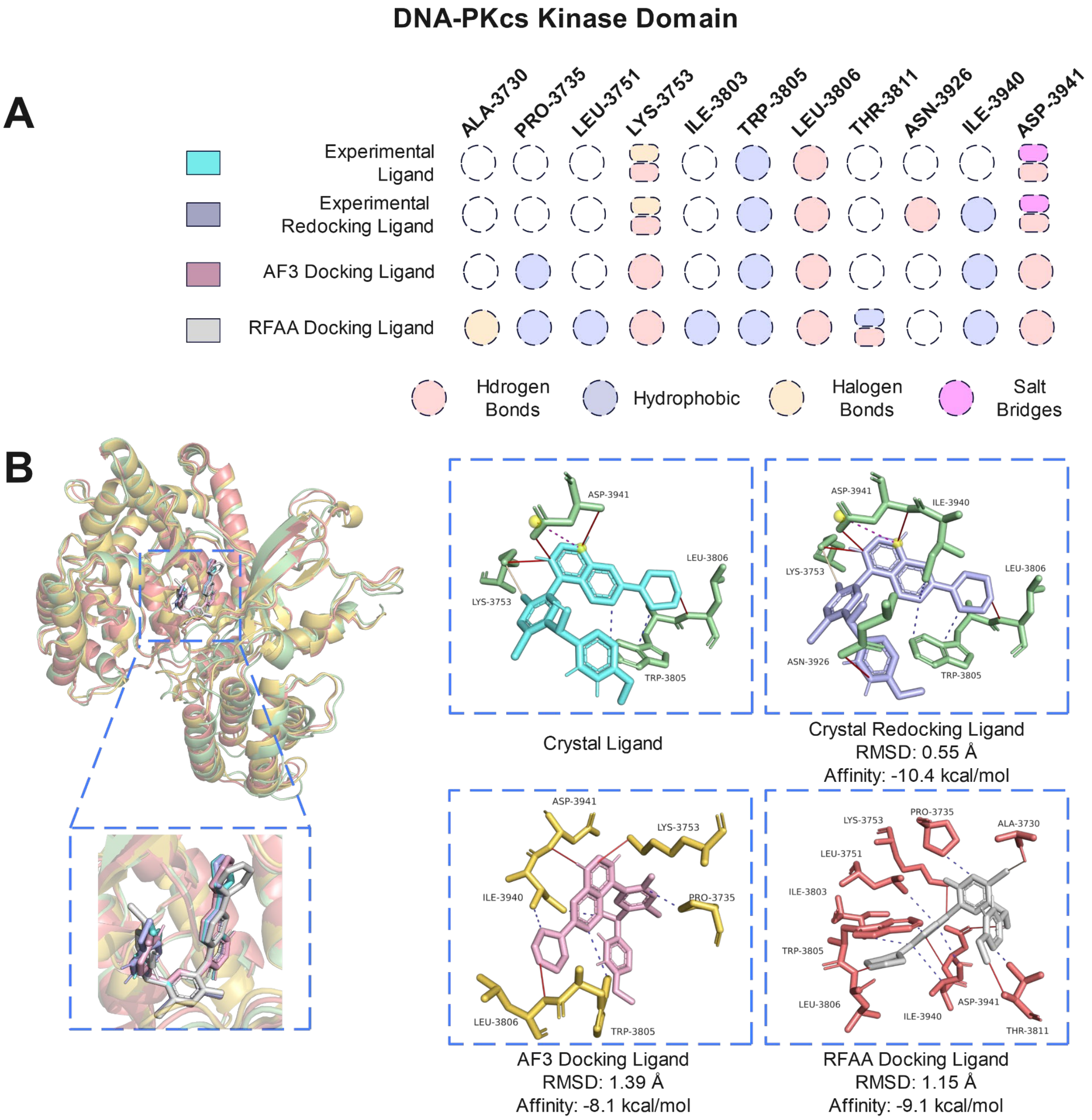


Figs. 4: The interaction analysis of ligand M3814 docked with the experimental structure, AF3, and RFAA model of DNA-PKcs. (A) The interactions between M3814 and DNA-PKcs KD structures from different sources. (B) Conformations of M3814 with experimental and AI-predicted DNA-PKcs KD structures, along with the RMSD values compared to the original structure and the affinity scores from molecular docking.

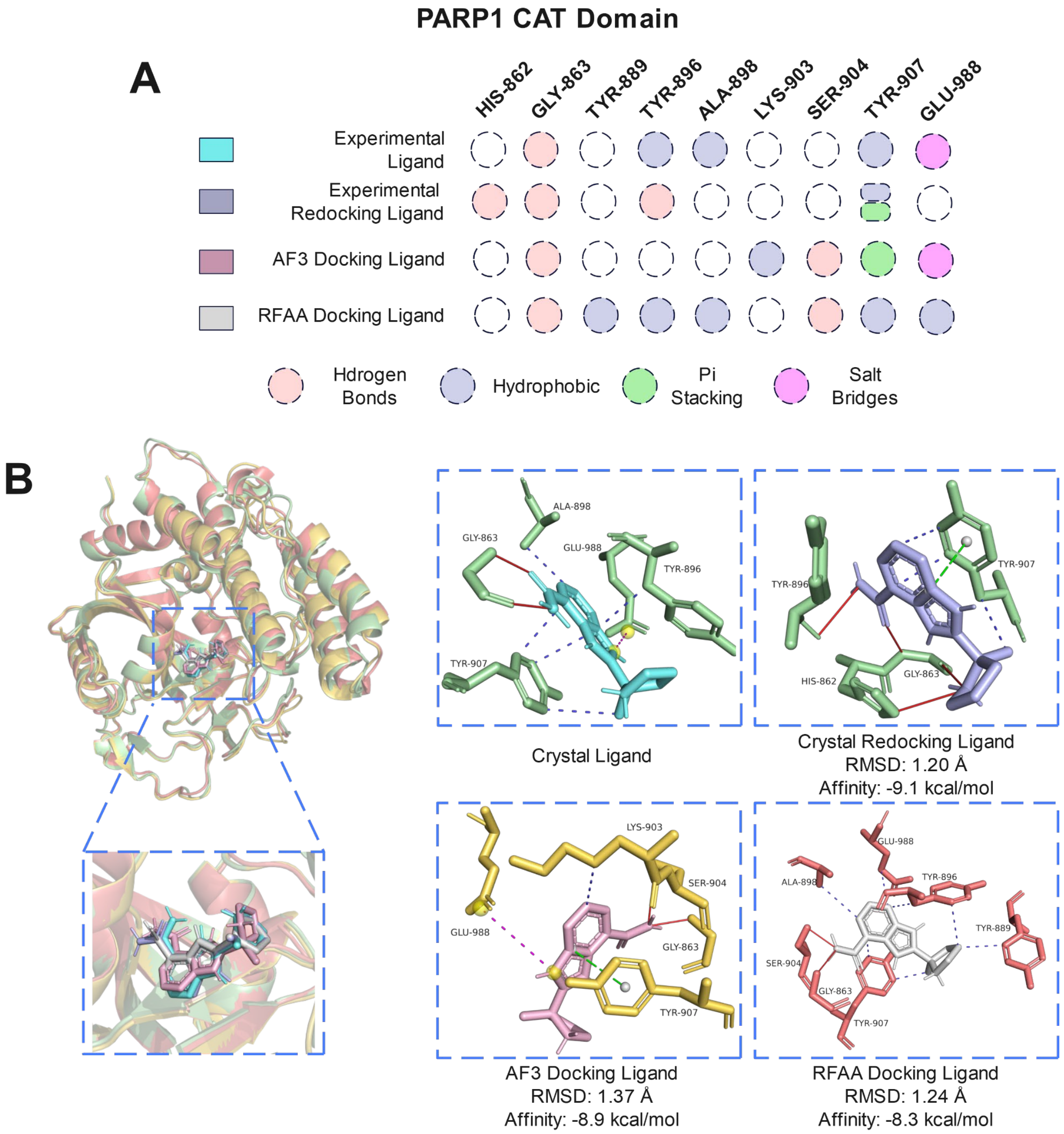


Figs. 5: The interaction analysis of ligand veliparib docked with the experimental structure, AF3, and RFAA model of PARP1. (A) The interactions between veliparib and PARP1 CAT structures from different sources. (B) Conformations of veliparib with experimental and AI-predicted PARP1 CAT structures, along with the RMSD values compared to the original structure and the affinity scores from molecular docking.

**AF3**

**Depth: 18.73Å Surface: 1026.58Å$^2$**

**Surface-Volume-Ratio: 1.34**

**Volume: 768.00Å$^3$**

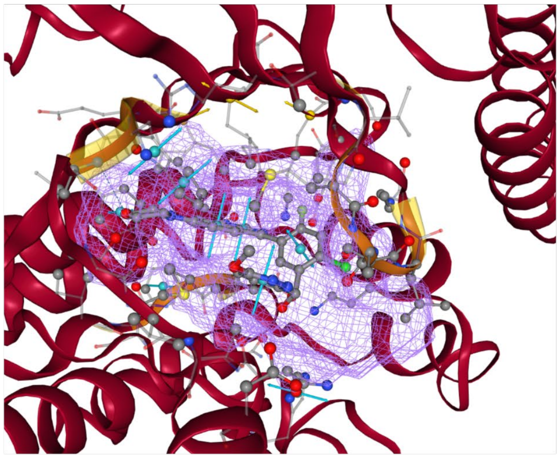

**AF2**

**Depth: 21.50Å Surface: 892.47Å$^2$**

**Surface-Volume-Ratio: 1.13**

**Volume: 792.58Å$^3$**

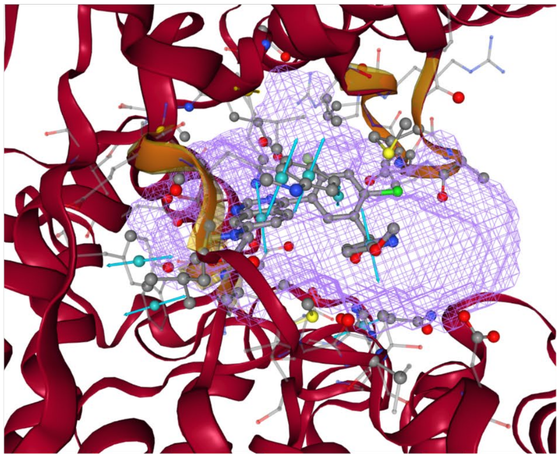

**Experimental**

**Depth: 19.30Å Surface: 984.12Å$^2$**

**Surface-Volume-Ratio: 1.00**

**Volume: 983.04Å$^3$**

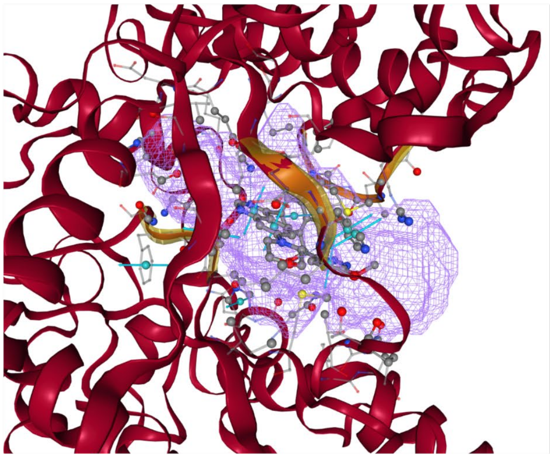

**RFAA**

**Depth: 20.71Å Surface: 921.33Å$^2$**

**Surface-Volume-Ratio: 1.06**

**Volume: 871.42Å$^3$**

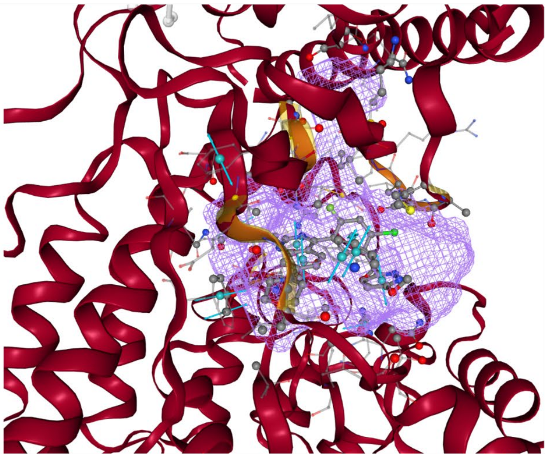

Figs. 6: The binding pockets parameters of DNA-PKcs structures. Four structures generated from experimental, AF3, AF2, RFAA. We focused on their differences on parameters of depth, surface, surface-volume-ratio(svr), and volume.

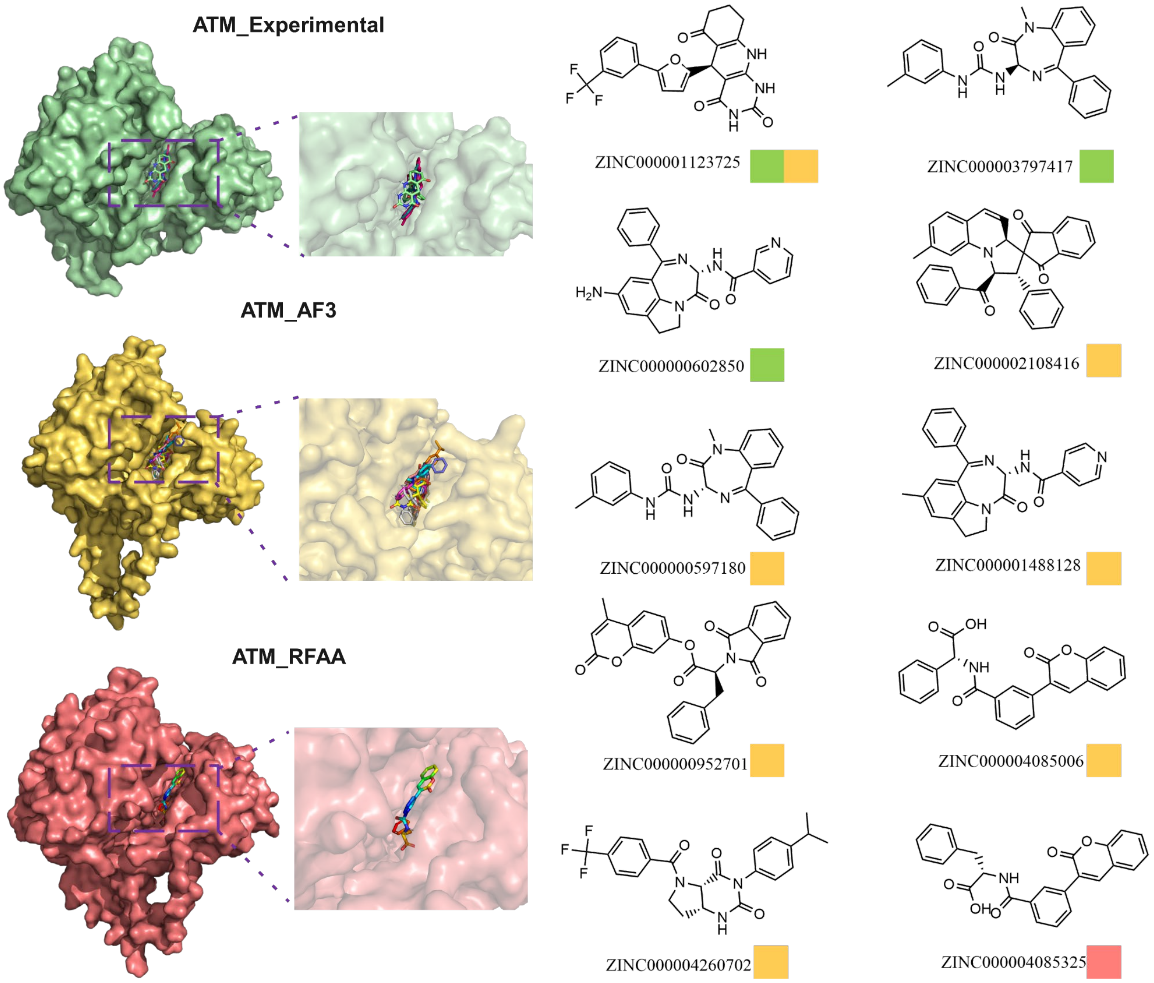


Figs. 7: The filtered compounds identified by ATM structures. Ten molecules (‘3725, ‘7417, ‘2850, ‘8416, ‘7180, ‘8128, ‘2701, ‘5006, ‘0702, and ‘5325) were identified as candidates by clustering top-ranked molecules, screening their druggability and toxicities, and merging molecules from different structural sources (Experimental, AF3, and RFAA). There are three molecules that were identified by the ATM_Experimental (Green), seven molecules were identified by ATM_AF3 (Yellow), one molecule was identified by ATM_RFAA (Red).

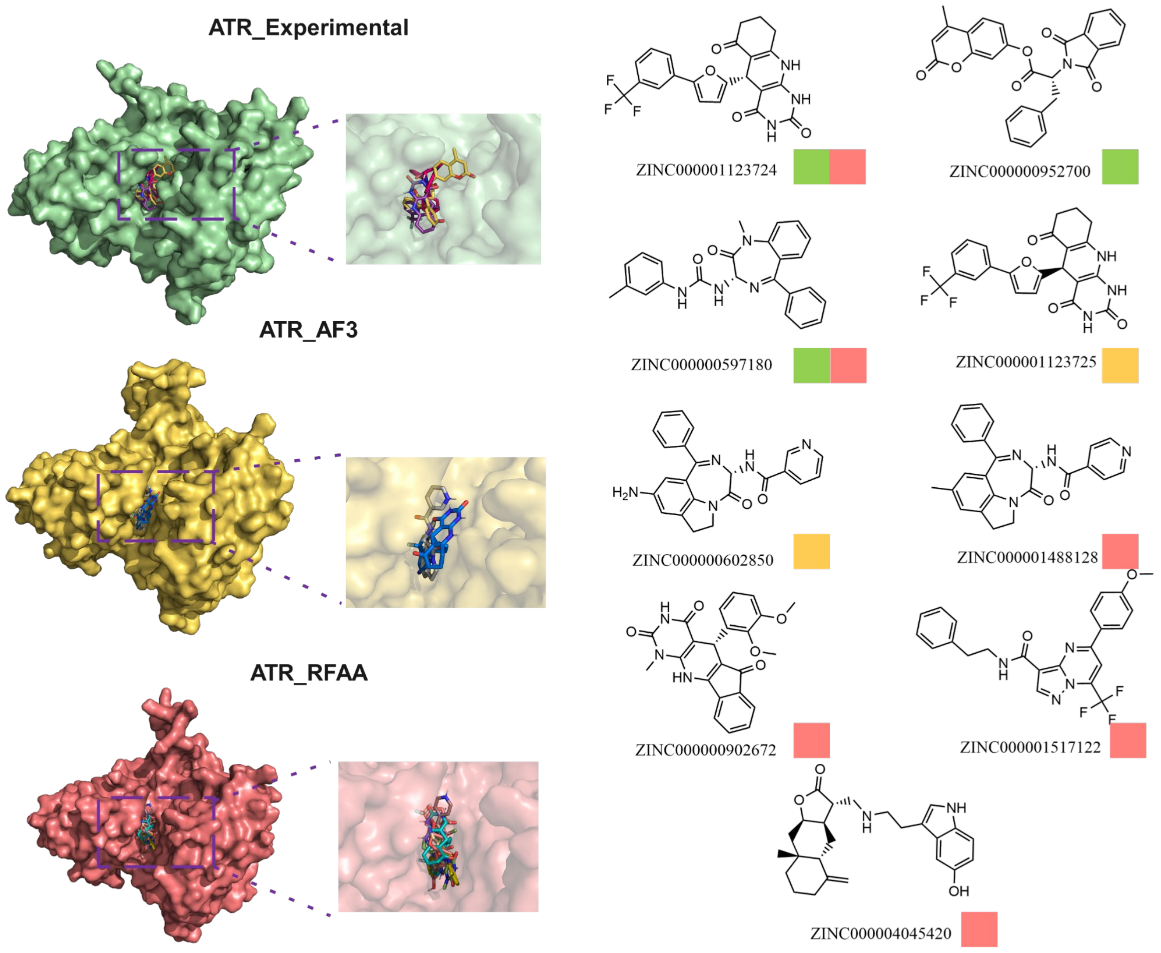


Figs. 8: The filtered compounds identified by ATR structures. Nine molecules (‘3724, ‘2700, ‘7180, ‘3725, ‘2850, ‘8128, ‘2672, ‘7122, and ‘5420) were identified as candidates by clustering top-ranked molecules, screening their druggability and toxicities, and merging molecules from different structural sources (Experimental, AF3, and RFAA). There are three molecules that were identified by the ATR_Experimental (Green), two molecules were identified by ATR_AF3 (Yellow), six molecules were identified by ATR_RFAA (Red).

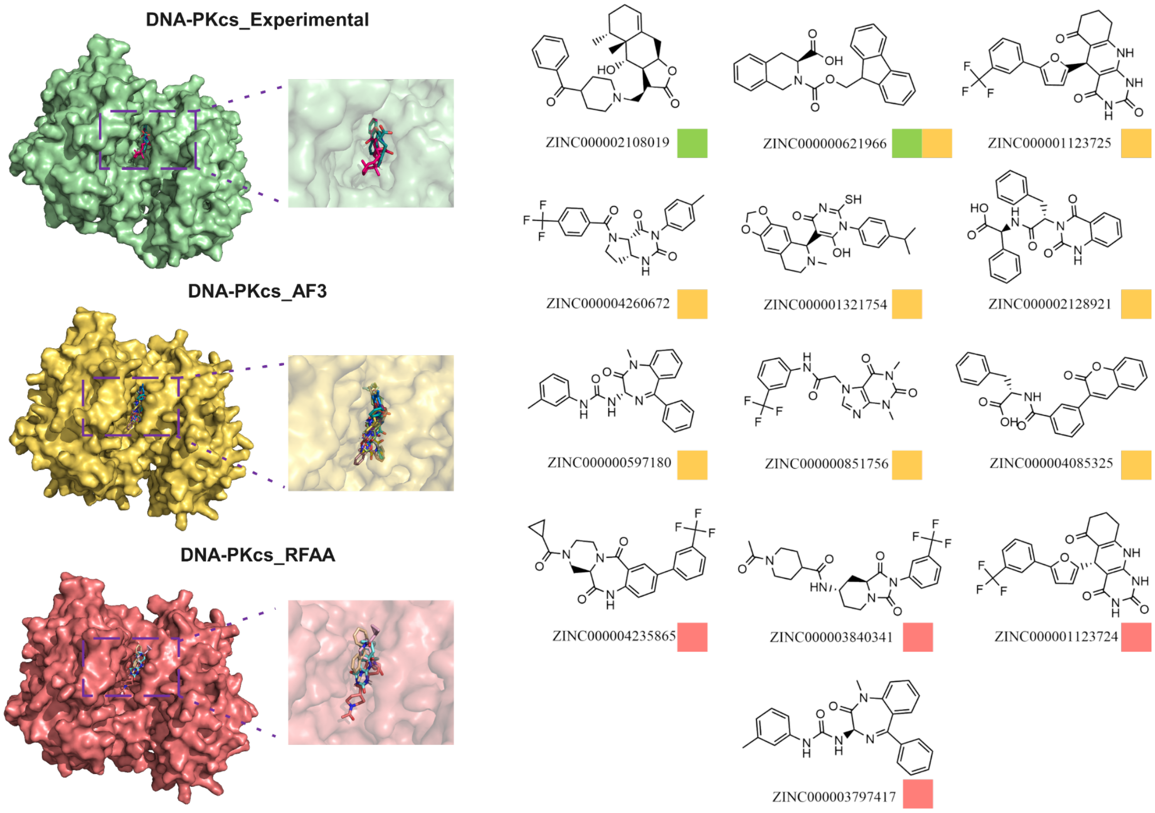


Figs. 9: The filtered compounds identified by DNA-PKcs structures. Thirteen molecules (‘8019, ‘1966, ‘3725, ‘0672, ‘1754, ‘8921, ‘7180, ‘1756, ‘5325, ‘5865, ‘0341, ‘3724, and ‘7417) were identified as candidates by clustering top-ranked molecules, screening their druggability and toxicities, and merging molecules from different structural sources (Experimental, AF3, and RFAA). There are two molecules that were identified by the DNA-PKcs_Experimental (Green), eight molecules were identified by DNA-PKcs_AF3 (Yellow), four molecules were identified by DNA-PKcs_RFAA (Red).

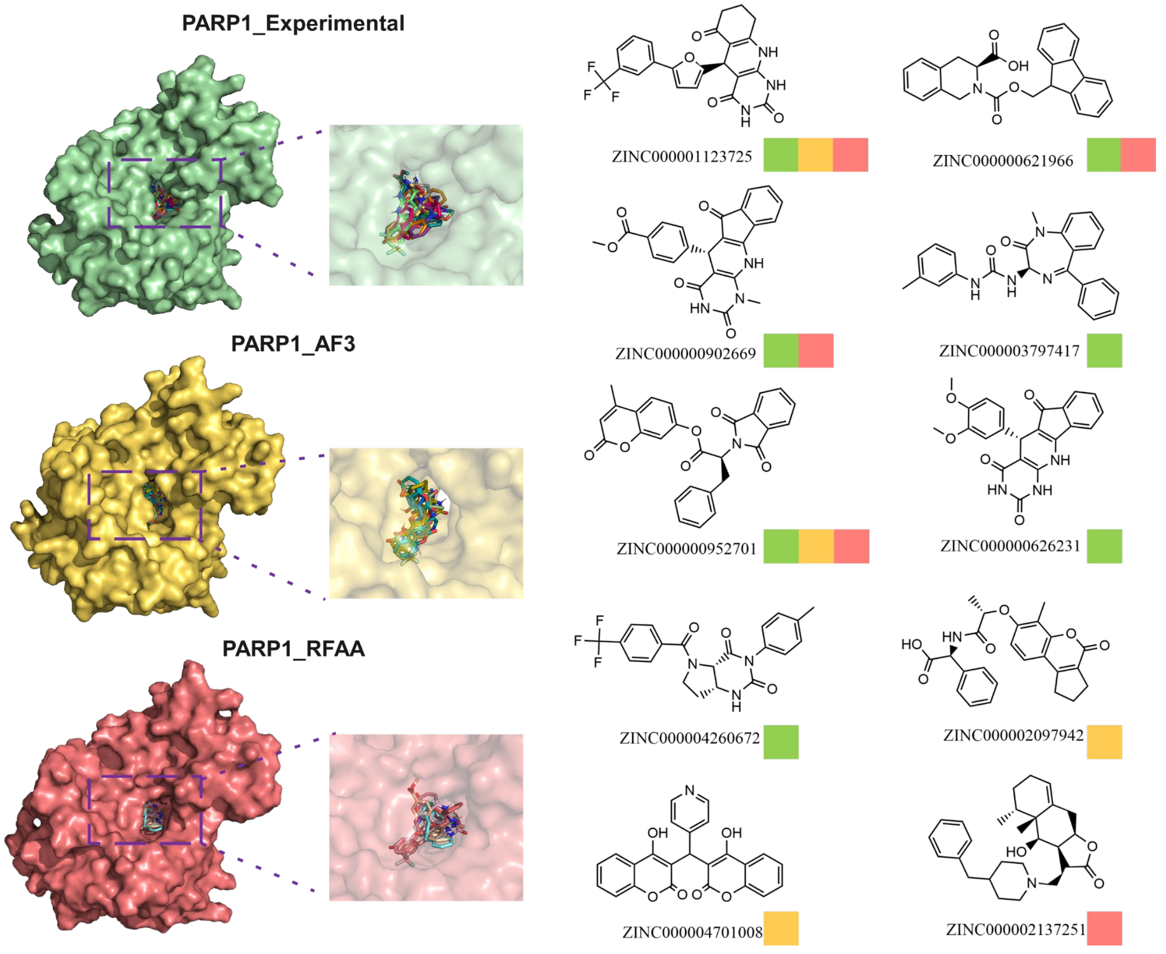


Figs. 10: The filtered compounds identified by PARP1 structures. Ten molecules (‘3725, ‘1966, ‘2669, ‘7417, ‘2701, ‘6231, ‘0672, ‘7942, ‘1008, and ‘7251) were identified as candidates by clustering top-ranked molecules, screening their druggability and toxicities, and merging molecules from different structural sources (Experimental, AF3, and RFAA). There are seven molecules identified by PARP1_Experimental (Green), four molecules were identified by PARP1_AF3 (Yellow), five molecules were identified by PARP1_RFAA (Red).

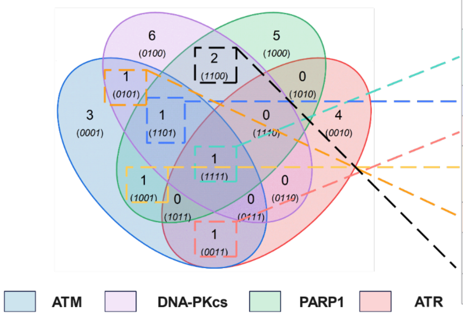


| ZINC IDs | Targets | ATM | ATR | DNAPKCS | PARP1 |
|---|---|---|---|---|---|
| ZINC000001123725 | ATM ATR DNAPKCS PARP1 | Crystal AF3 | AF3 | AF3 | Crystal AF3 RFAA |
| ZINC000003797417 | ATM DNAPKCS PARP1 | Crystal | | RFAA | Crystal |
| ZINC000000602850 | ATM ATR | Crystal | AF3 | | |
| ZINC000000952701 | ATM PARP1 | AF3 | | | Crystal AF3 RFAA |
| ZINC000004085325 | ATM DNAPKCS | RFAA | | AF3 | |
| ZINC000000621966 | DNAPKCS PARP1 | | | Crystal AF3 | Crystal RFAA |
| ZINC000004260672 | DNAPKCS PARP1 | | | AF3 | Crystal |

Figs. 11: Seven potential multi-target compounds. ZINC000001123725 ('3725), ZINC000003797417 ('7417), ZINC000000602850 ('2850), ZINC000000952701 ('2701), ZINC000004085325 ('5325), ZINC000000621966 ('1966), ZINC000004260672 ('0672) were identified by calculating the intersections of candidate lists of ATM, ATR, DNA-PKcs, and PARP1.

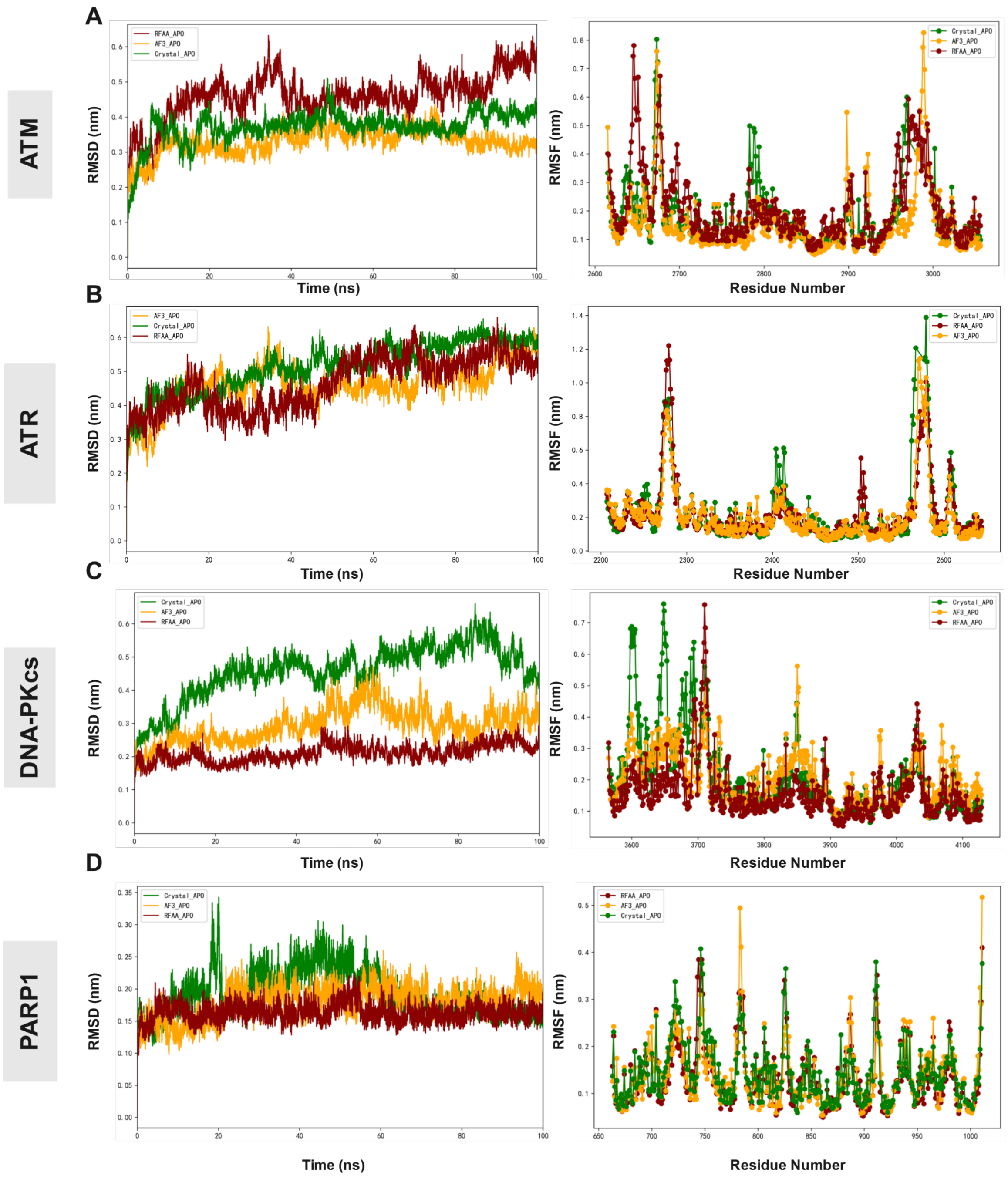


Figs. 12: The molecular dynamics simulations (MDS) of Apo forms of 12 protein structures. (A) The Apo forms of experimental (Green), AF3 (Yellow), and RFAA (Red) structures for ATM protein were simulated for 100ns. The RMSD reflected the backbone stability, while the RMSF showed the flexibility of residues located in its kinase domains. (B) The 100ns MDS on Apo forms of ATR protein. (C) The 100ns MDS on Apo forms of DNA-PKcs protein. (B) The 100ns MDS on Apo forms of PARP1 protein.

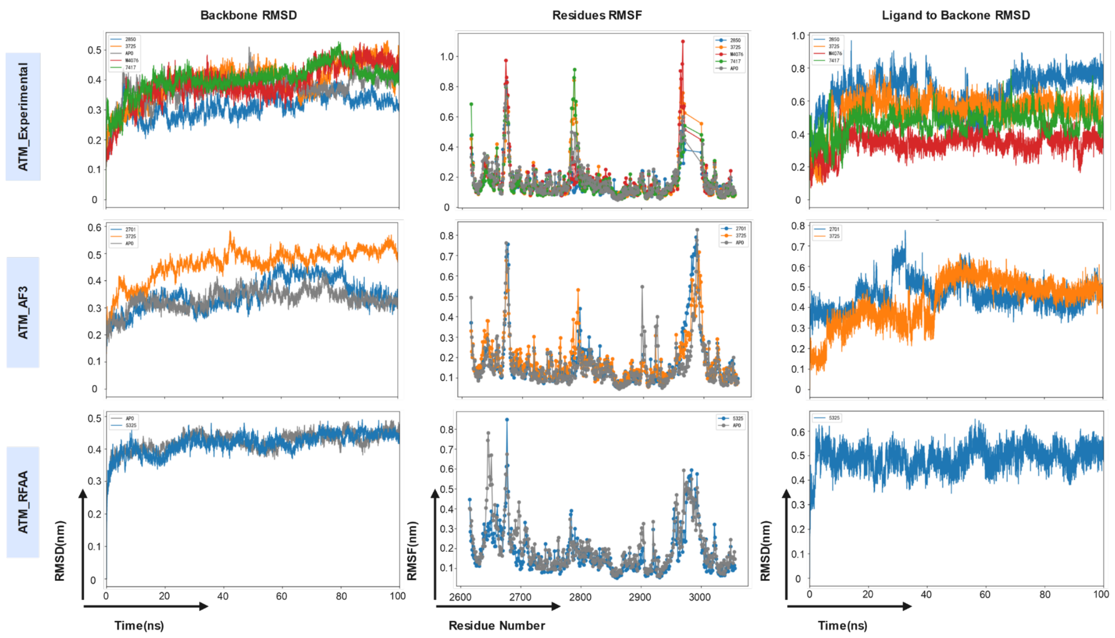


Figs. 13: The 100ns molecular dynamics simulations (MDS) of protein-ligand assemblies screened by ATM protein. Five ligands ('2850, '3725, '7417, '2701, and '5325) and the reference ligand, M4076, were formed assemblies with different ATM structures. The RMSD of protein backbones were compared to the baselines of Apo forms (Gray). Residues in the kinase domains of ATM protein were calculated for their RMSF for flexibility evaluation. The RMSD of ligand relative to protein backbone was used to evaluate the convergence of the simulation systems.

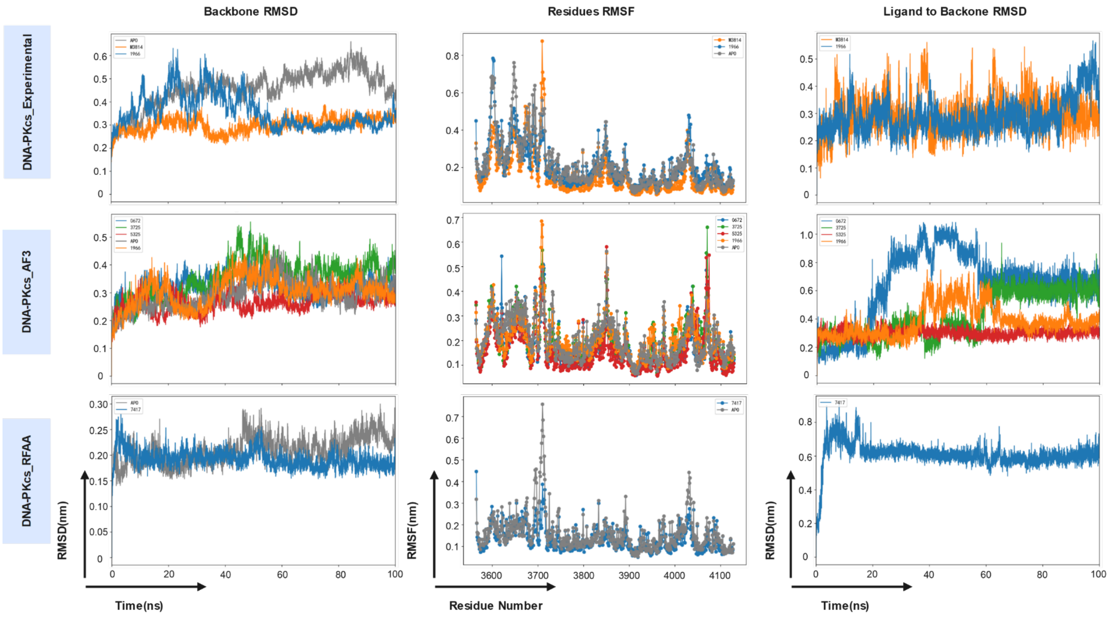


Figs. 14: The 100ns molecular dynamics simulations (MDS) of protein-ligand assemblies screened by DNA-PKcs protein. Five ligands ('1966, '0672, '3725, '5325, and '7417) and the reference ligand, M3814, were formed assemblies with different DNA-PKcs structures. The RMSD of protein backbones were compared to the baselines of Apo forms (Gray). Residues in the kinase domains of DNA-PKcs protein were calculated for their RMSF for flexibility evaluation. The RMSD of ligand relative to protein backbone was used to evaluate the convergence of the simulation systems.

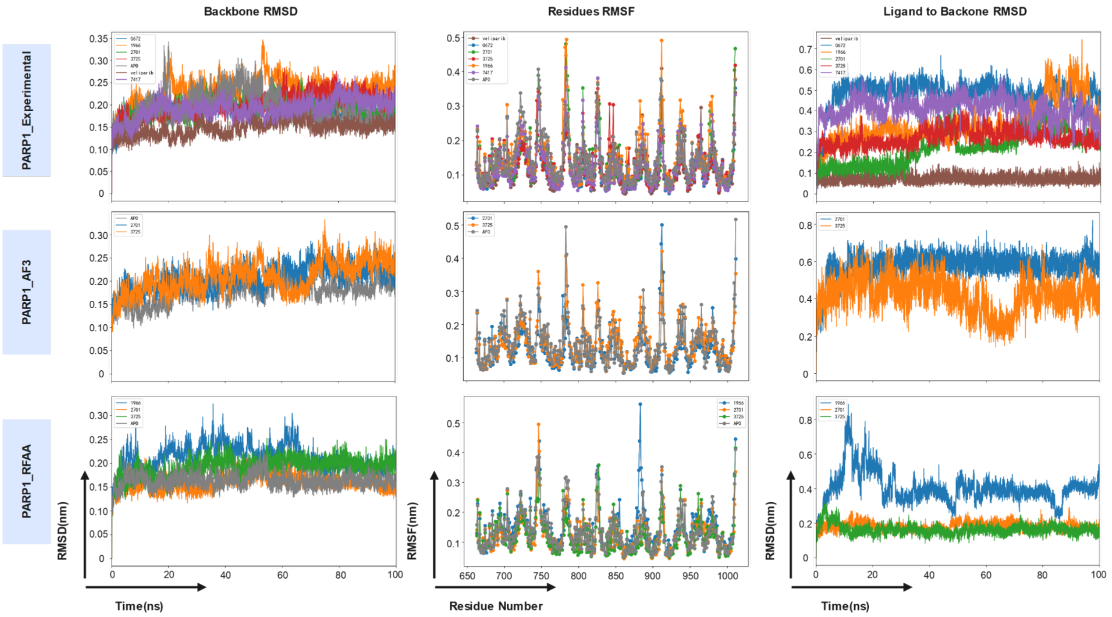


Figs. 15: The 100ns molecular dynamics simulations (MDS) of protein-ligand assemblies screened by PARP1 protein. Five ligands ('0672, '1966, '2701, '3725, and '7417) and the reference ligand, veliparib, were formed assemblies with different PARP1 structures. The RMSD of protein backbones were compared to the baselines of Apo forms (Gray). Residues in the CAT domains of PARP1 protein were calculated for their RMSF for flexibility evaluation. The RMSD of ligand relative to protein backbone was used to evaluate the convergence of the simulation systems.

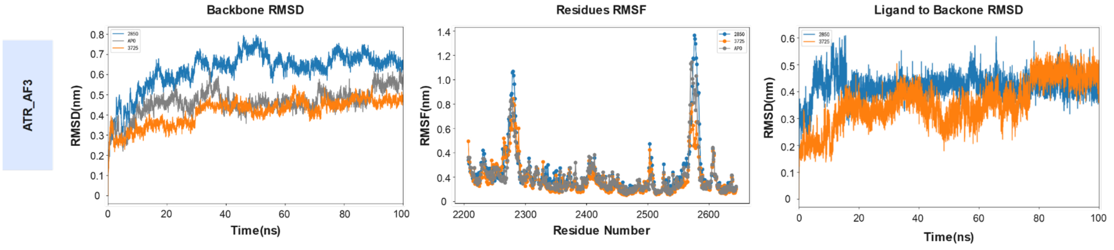


Figs. 16: The 100ns molecular dynamics simulations (MDS) of protein-ligand assemblies screened by ATR structure. Two ligands ('2850 and '3725) were formed assemblies with ATR_AF3 structure. The RMSD of protein backbones were compared to the baselines of Apo forms (Gray). Residues in the kinase domains of ATR_AF3 structure were calculated for their RMSF for flexibility evaluation. The RMSD of ligand relative to protein backbone were used to evaluate the convergence of the simulation systems

Table s1. Changes in the number of compounds in target structures with different filtering criteria. Top-ranked compounds were clustered using their morgan fingerprints, with Tanimoto coefficient (Tc) = 0.5. The compound with the highest docking score was selected as representative for the following druggability, toxicity, and novelty screening. These filtered compounds were regarded as hit compounds. Hit compounds identified by different sourced structures were merged as total hit compounds. Ultimately, seven promising multi-target compounds identified by simultaneously in at least two targets were selected.

| **Target Structures** | | **Top-ranked compounds** | **Represent compounds** | **Filtered Hit compounds** | **Total Hit compounds** | **Multi-target compounds** |
|---|---|---|---|---|---|---|
| **ATM** | Experimental | 1176 | 250 | 3 | 10 | 7 |
| | AF3 | 1090 | 270 | 7 | | |
| | RFAA | 1008 | 154 | 1 | | |
| **ATR** | Experimental | 1075 | 211 | 3 | 9 | |
| | AF3 | 1162 | 277 | 2 | | |
| | RFAA | 1109 | 264 | 6 | | |
| **DNA-PKcs** | Experimental | 1250 | 196 | 2 | 13 | |
| | AF3 | 1066 | 237 | 8 | | |
| | RFAA | 1142 | 184 | 4 | | |
| **PARP 1** | Experimental | 1058 | 217 | 7 | 10 | |
| | AF3 | 1143 | 267 | 4 | | |
| | RFAA | 1249 | 275 | 5 | | |
